# Structural Requirements for Reverse Transcription by a Diversity-generating Retroelement

**DOI:** 10.1101/2023.10.23.563531

**Authors:** Sumit Handa, Tapan Biswas, Jeet Chakraborty, Gourisankar Ghosh, Blair G. Paul, Partho Ghosh

## Abstract

Diversity-generating retroelements (DGRs) create massive protein sequence variation in ecologically diverse microbes. Variation occurs during reverse transcription of a protein-encoding RNA template coupled to misincorporation at adenosines. In the prototypical *Bordetella* bacteriophage DGR, the template must be surrounded by upstream and downstream RNA segments for cDNA synthesis by the reverse transcriptase bRT and associated protein Avd. The function of the surrounding RNA was unknown. Cryo-EM revealed that this RNA enveloped bRT and lay over barrel-shaped Avd, forming an intimate ribonucleoprotein (RNP).

An abundance of essential interactions between RNA structural elements and bRT-Avd precisely positioned an RNA homoduplex for initiation of cDNA synthesis by *cis*-priming. Our results explain how the surrounding RNA primes cDNA synthesis, promotes processivity, terminates polymerization, and strictly limits mutagenesis to select proteins through mechanisms that are likely conserved in DGRs from distant taxa.

## Introduction

Diversity-generating retroelements (DGRs) create massive protein sequence variation (up to 10^30^) in ecologically diverse bacteria and archaea, and their viruses. A recent survey identified ∼31,000 DGRs from > 1,500 bacterial and archaeal genera, constituting > 90 environment types (*1*). DGRs are especially enriched in the human gut microbiome (*1, 2*) and nano-sized microbes, which appear to comprise a major fraction of microbial life. The latter maintain DGRs despite having reduced genomes (*3, 4*). DGRs are also implicated in the emergence of multicellularity (*5, 6*). The prototypical DGR of *Bordetella* bacteriophage (BΦ) can produce ∼10^13^ variants of its receptor-binding protein Mtd (*7, 8*), which enables the BΦ to use alternative receptors when its usual receptor is missing due to gene expression changes in *Bordetella*. This rapid adaptation to a dynamic environment is likely the general function of DGRs.

The BΦ DGR consists of three genes and two repeats (Fig. 1a). The genes encode the variable protein Mtd, a reverse transcriptase (bRT), and an accessory variability determinant (Avd) protein (*7*). The repeats are the variable region (*VR*), which encodes the variable amino acids of Mtd, and the template region (*TR*), which is nearly identical to *VR* except mainly at adenosines. Sequence variation arises during reverse transcription of an RNA copy of *TR* through frequent misincorporation at *TR* adenosines. *TR-*cDNA, containing errors at positions corresponding to adenosines, then homes to and replaces the extant copy of *VR* to give rise to an Mtd variant. The mechanism of homing is poorly understood. *TR* is sequence invariant and thus there are no limits on the number of rounds of adenosine (A)-mutagenesis that can occur.

**Fig. 1.**
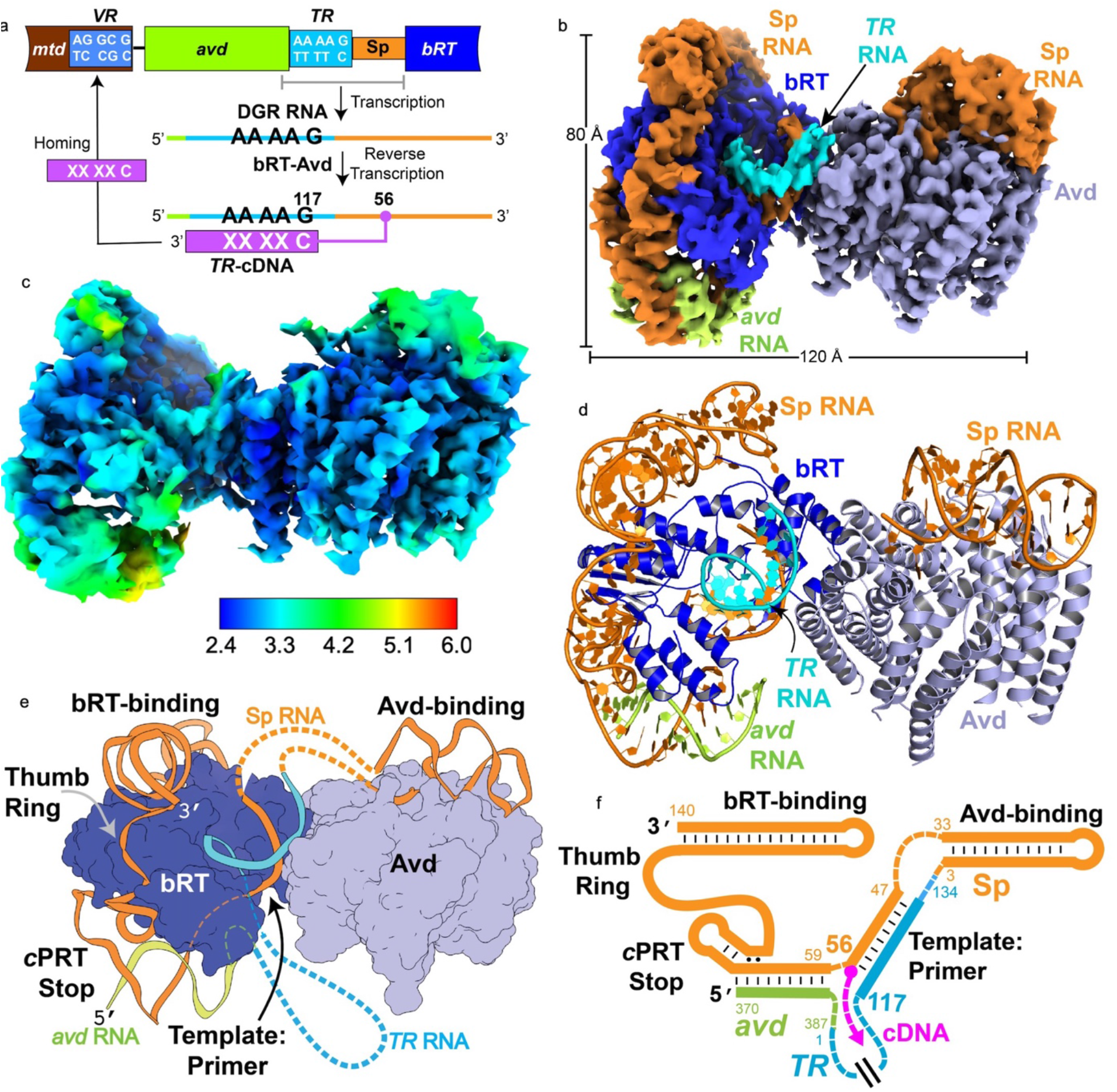
DGR and bRT-Avd:RNA^Δ98^ complex. **a**. Schematic ofB<I> DGR. *VR* and *TR* (both strands depicted) differ mainly at adenosines in *TR* (top strand). An example sequence is shown for *VR*, in which three of the four adenosines in *TR* were mutated in a previous round of A-mutagenesis. ‘X’ denotes misincorporation errors at adenosines. Thin lines correspond to RNA, and boxes to DNA. **b-d** Cryo-EM density (b), local resolution map with resolution scale in Å below (c), and cartoon (d) ofbRT-Avd:RNA^Δ98^ (bRT, blue; Avd, slate; RNA^Δ98^ Iight green for *avd, cyan* for TR, orange for Sp). **e and f**. Schematic of RNA elements in three-dimensional (e) and linear representation (f). Dotted lines indicate sections that were either too flexible to model (parts of *avd, TR*, or Sp) or were absent in the complex (TRI 0-107 and cDNA).

Reverse transcription is carried out by a complex of bRT and Avd. Unlike other RTs, bRT lacks reverse transcription activity on its own and requires association with Avd (*9*). bRT belongs to a monophyletic clade of DGR reverse transcriptases (RTs) most closely related to group II intron RTs (*1, 10, 11*), and Avd forms a pentameric, positively-charged barrel-like structure similar to S23 ribosomal proteins (*12*). Avd-like proteins occur in the great majority of DGRs (*13*). The bRT-Avd complex has intrinsic A-mutagenesis activity on RNA or DNA templates (*9, 14*).

Significantly, *TR* RNA is insufficient for reverse transcription by bRT-Avd and must instead be surrounded by *cis-*acting elements (*9*). Most prominent is a 140-nucleotide (nt) noncoding RNA segment called Spacer (Sp) located immediately downstream of *TR* (Fig. 1a). Sp RNA supplies the priming site (^Sp^A56) for cDNA synthesis (*9, 15*). The 2’-OH of ^Sp^A56 is necessary for cDNA synthesis (*9*), suggesting that it acts as the priming nucleophile, as occurs in retrons (*16*). Thus, reverse transcription is *cis*-primed and a covalently linked RNA-cDNA molecule is produced. The first nucleotide reverse transcribed is ^*TR*^G117 (*9, 15*). A 20 nt region upstream of *TR*, consisting of the 3’-terminal codons of *avd*, is also required for *cis-*primed reverse transcription (*c*PRT). We use the term ‘DGR RNA’ to refer to the longer RNA required for *c*PRT, containing 20 *avd* and 140 Sp nts surrounding *TR*. This longer DGR RNA confers not only *cis-*priming but also processive polymerization, resulting in ∼120 nt cDNAs that span *TR* (*9*), in contrast to the 5-30 nts cDNAs synthesized from non-cognate RNA templates using exogenous primers (*9*).

### Structure Determination

How the DGR RNA combines with bRT-Avd for cDNA synthesis was unknown and pursued through cryo-EM. Initial maps of the bRT-Avd:RNA complex were limited to ∼7 Å resolution due to RNA flexibility and preferred particle orientation. To limit RNA flexibility, the central 98 nts of *TR* (Δ^*TR*^10-107) were deleted from DGR RNA (Fig. S1a). *TR* has an informational rather than functional role (*17*), and the truncated RNA (called RNA^Δ98^) has all the essential *cis-*acting elements. Preferred orientation was unexpectedly alleviated by fusing pentamer-forming *Methylobacterium extorquens* formaldehyde activating enzyme (Fae) to the Avd C-terminus, in an attempt to increase the mass of the ∼176 kDa bRT-Avd:RNA^Δ98^ complex. The bRT-(Avd-Fae):RNA^Δ98^ (∼258 kDa) complex yielded maps that extended to as high as ∼3.1 Å resolution (Tables S1 and S2, Figs. S2-S5). Fusions of other pentamer-forming proteins did not yield a similar improvement. bRT-(Avd-Fae) reverse transcribed RNA^Δ98^ and misincorporated at adenosine at the same frequency as bRT-Avd (Figs. S1b and c) (*14*). Fae was positionally variable and masked out, and thus we refer to Avd rather than Avd-Fae hereafter.

The most interpretable map came from a complex that contained RNA^Δ98^ with a ^*TR*^U114A substitution and had been incubated with dCTP, dATP, and ddGTP (Figs. 1b-d, Table S1, Complex 1). The intent was to visualize a short cDNA strand and ^*TR*^U114A at the template site, but the population of such complexes was too sparse for class-averaging, consistent with < 2% of templates yielding cDNA (*14*). Instead, this sample yielded a complex that had not yet commenced synthesis. Most of bRT and Avd were in density with a local resolution better than the global 3.25 Å resolution (Figs. 1c and S6a, b), and the entirety of bRT and Avd, except for a few N- and C-terminal amino acids in the latter, were modeled with an excellent fit (Table S2) (*18*). Segments of RNA that contacted bRT or Avd were generally in < 3.25 Å resolution density (Figs. 1c and 6b), while segments not in contact were often in ∼4-5 Å resolution density. In total, the RNA also had an excellent fit to the density (Table S2), and while there were a few breaks in the chain, ∼75% could be modeled.

### Intimate Ribonucleoprotein

Most strikingly, the structure revealed an intimate ribonucleoprotein (RNP) in which bRT was almost entirely enveloped by Sp RNA, which also lay over the narrow end of the Avd barrel (Fig. 1). In between, Sp RNA formed two adjacent duplexes, one with *TR* and the other with the upstream *avd* segment. bRT most closely resembled group II intron RTs as predicted (*11, 19-21*) (Figs. 2a and S6c), and shared with these proteins and non-LTR RTs (*22*) an N-terminal extension subdomain (NTE). Following the NTE were the canonical finger, palm, and thumb subdomains of nucleotide polymerases. bRT lacked additional subdomains found in related RTs, such as DNA-binding, RNA-binding, or endonuclease. The bRT NTE contacted Avd (two of five protomers) through an extensive and mainly electrostatic interface. Bound and free Avd were structurally similar (1.4 Å rmsd, 456 Cα), and the bRT-Avd interface was consistent with prior mutagenesis results (*12*) (Fig. S6d). While Avd is required for catalysis (*9*), its closest approach to the bRT active site was ∼40 Å, suggesting that it is not directly involved in chemistry.

**Fig. 2.**
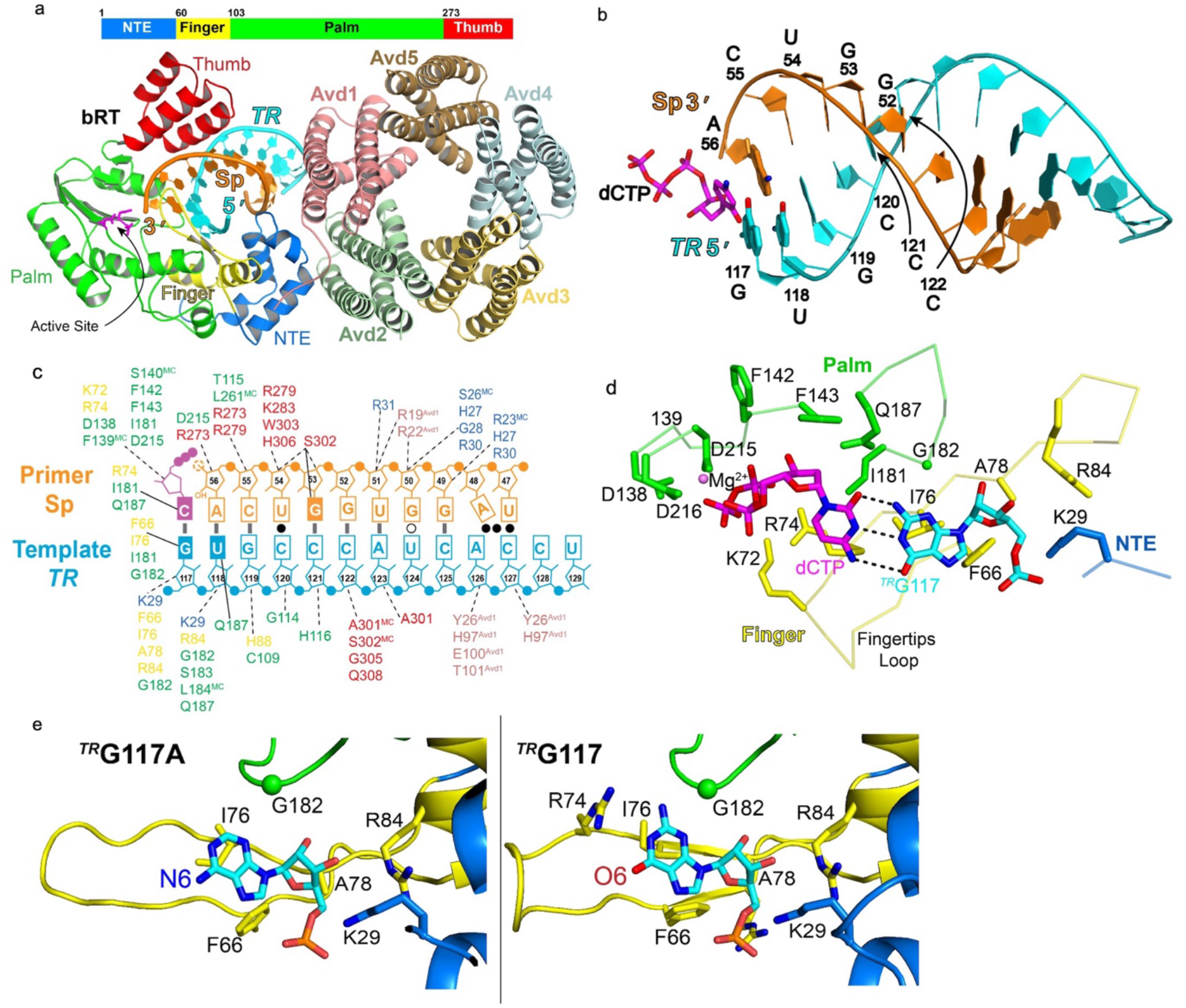
bRT-Avd and Template:Primer Duplex. **a**. bRT (schematic and structure: NTE, blue; finger, yellow; palm, green; thumb, red), Avd (colored individually), and *TR*:Sp template:primer duplex (cyan and orange, respectively). bRT active site amino acids Dl38, D215, and D216 in magenta. **b**. *TR*:Sp template:primer duplex with dCTP in magenta. **c**. Schematic of *TR*:Sp template:primer duplex with base pairs shown (solid bar, Watson-Crick; open circle, wobble; and closed circle, non-standard). Base-specific (sold lines) and ribose or phosphate (dotted lines) contacts from bRT or Avd shown. Colors oflabels correspond to coloring in panel a. The side chains ofbRT Ql87 and S301 are 3.9 Å from the 02 of^*TR*^(Jl 18 and 3.4 Å from the N3 of^Sp^53G, respectively. **d**. Interactions between ^*TR*^G 117 (carbon and phosphorus, cyan) at the T-site and dCTP (carbon and phosphorus, magenta) at the N-site (for both nucleotides, nitrogen blue, and oxygen red), and with bRT amino acids (colored as in panel a). Hydrogen bonds shown as black dotted lines. The putative Mg^2+^was 2.2 and 2.3 Å from Dl38 and D215, respectively, and 3.3 Å from one of the oxygens of the a-phosphate. **e**. Interactions between (left) ^*TR*^G 117 A and (right) ^*TR*^G 117 (carbon and phosphorus, cyan) at the T-site with bRT (carbon colored by subdomain, as in panel a; and oxygen red, and nitrogen blue for nucleotides and protein).

### Template:Primer Duplex in Active Site

The bRT active site was occupied by an RNA homoduplex in which one strand (^*TR*^117-129) contained the initiating nucleotide ^*TR*^G117, and the other (^Sp^47-56) the priming nucleotide ^Sp^A56 (Figs. 2 and S7). The bRT finger, palm, and thumb contacted the end of the template:primer duplex proximal to the active site, while the NTE and an Avd protomer contacted the distal end. This suggested that the role of Avd in catalysis is to enforce proper positioning of the template:primer duplex either through direct contact or indirectly through the NTE subdomain. The *TR*:Sp duplex formed seven canonical base pairs (bps), as predicted (*9, 15*), along with three non-standard pairs (Figs. 2b, c) (*9, 15*). The four base pairs closest to the active site, spanning ^Sp^53-56 and ^*TR*^118-121, are functionally required whereas the others are not (*9, 15*). Bases of the *TR*:Sp RNA homoduplex were not contacted closely by bRT or Avd, the exception being the template base ^*TR*^G117 (Figs. 2c and d), which was contacted by conserved fingertips loop (F66 and I76) and palm (I181 and G182) amino acids. The NTE subdomain contacted the 5’ phosphate of ^*TR*^G117 through conserved Lys 29, which was on a loop braced by Avd (Fig. S7e). dCTP was at the incoming dNTP (N)-site, with cytosine forming canonical hydrogen bonds with the guanine at the template (T)-site. The cytosine was contacted by conserved fingertips loop (R74) and palm (I181 and Q187) amino acids, and stacked coaxially against the base of the priming nucleotide ^Sp^A56 (Fig. 2b). The 2’-OH of ^Sp^A56 could not be placed with chemical precision due to the resolution of the map, compounded by the lack of density for the following nucleotide, which would otherwise constrain its 3’-OH. The catalytic bRT aspartates 138, 215, and 216, and a cation, presumably Mg^2+^, were positioned adjacent to the dCTP phosphates (Fig. 2d). Thus, this complex appeared poised for catalysis and we termed it the ‘Active G:dCTP’ conformation (Table S1, Complex 1).

As bRT and other DGR RTs misincorporate specifically at template adenosines, we determined the 3.23 Å global resolution structure of an RNP that had a ^*TR*^G117A substitution in RNA^Δ98^. In this RNP, the adenine at ^*TR*^117 occupied the T-site, and while the nonhydrolyzable dCTP analog dCpCpp had been incubated with it, only broken density was observed at the N-site (Figs. 2e and S8, and Table S1, Complex 2). This RNP closely resembled one determined at 3.07 Å global resolution in which the T-site was occupied by guanine (^*TR*^G117 in wild-type RNA^Δ98^) and, despite incubation with dCpCpp, the N-site was similarly empty (Figs 2e and S8, Table S1, Complex 3). These two RNPs had an Active conformation (termed Active A:Empty and Active G:Empty, respectively), but differed from the Active G:dCTP conformation in having greater conformational variability. This was evident in the bRT finger and thumb subdomains and in the RNA. The *TR*:Sp template:primer duplex in these Active N-empty conformations, either with adenine or guanine at the T-site, occupied density of moderate local resolution (∼4 Å), in contrast to the higher local resolution in the Active G:dCTP conformation (Fig. S7). These observations suggest that the Active conformation does not absolutely depend on N-site occupancy but does seem to be stabilized by the correct incoming dNTP. Comparison of the N-empty complexes showed similar contacts by bRT to adenine and guanine at the T-site (Fig. 2e). Notably, neither bRT nor Avd contacted the substituent at the C6 position of these purines, which is a major determinant of A-mutagenesis (*14*).

### *c*PRT Stop

The template:primer duplex was immediately adjacent to a second duplex (Figs. 1e and f) in which one strand consisted of ^Sp^59-70 and the other the essential upstream *avd* segment (^*avd*^369-379) (Figs. 3 and S9). These strands formed 11 canonical bps, with ^Sp^U63 unpaired but accommodated within the coaxial stack. The *avd*:Sp duplex appeared to be part of a larger structural assembly, with the Sp strand continuing into a short stem-loop (^Sp^71-78) followed by three unpaired nucleotides (^Sp^79-81). bRT formed sparse contacts with the *avd*:Sp duplex, reflected in the moderate local resolution (∼4 Å) of its density, but numerous contacts to the short stem-loop and unpaired bases, which were in density of < 3 Å local resolution (Figs. 3b and c, S9). Two of the unpaired nucleotides (^Sp^A80 and A81) were inserted into the minor groove of G:C base pairs of the *avd*:Sp duplex, forming A-minor motifs and engaging in one of the few tertiary contacts in the RNA (Figs. 3d and e). The A-minor motif is one of the most prevalent tertiary interactions in RNA (*23, 24*). As observed here, it preferentially involves G:C base pairs, and often occurs as pairs or triples of adenosines. The unpaired adenosines were also contacted by bRT.

**Fig. 3.**
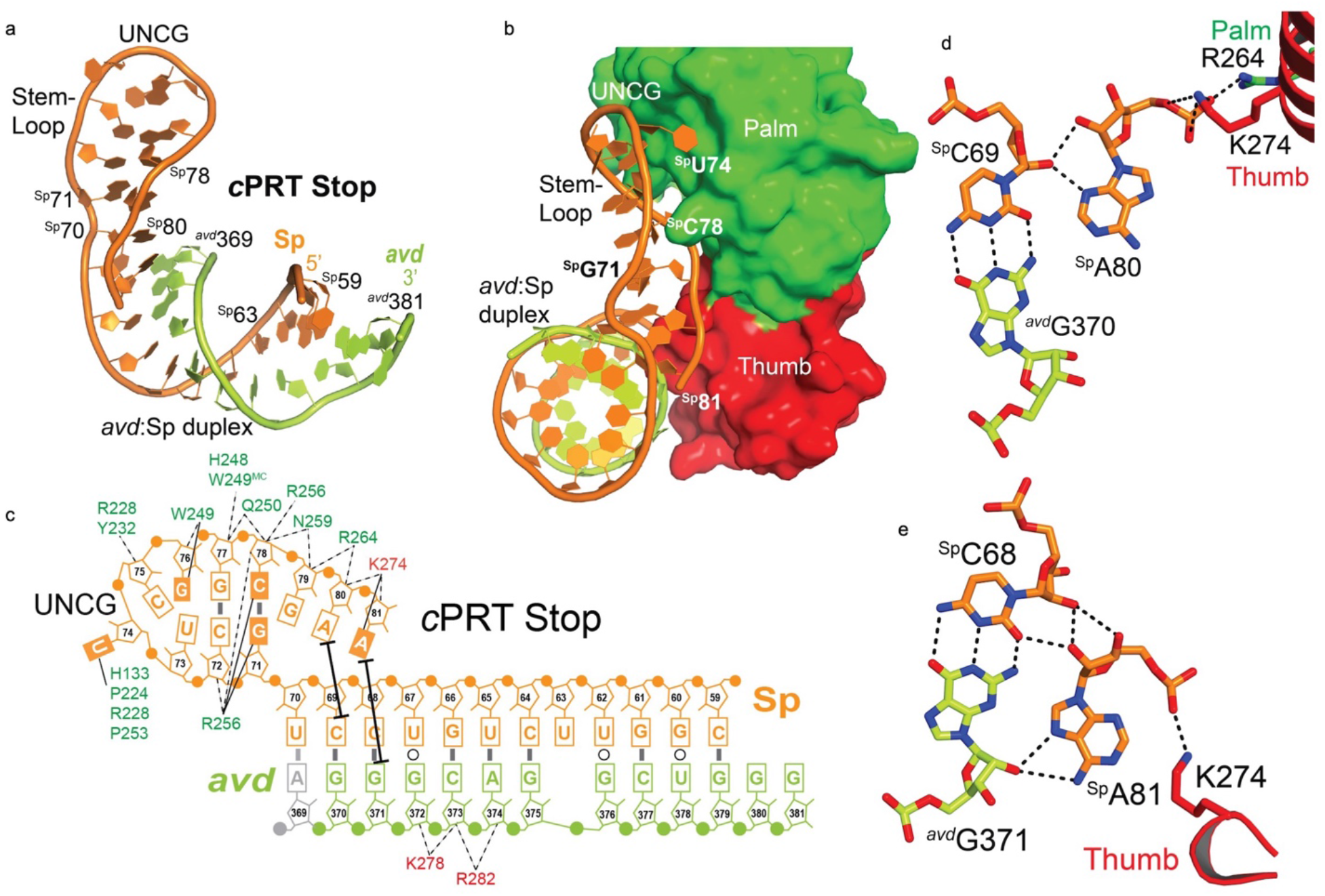
*c*PRT Stop. **a**. cPRT Stop (*avd*:Sp duplex, stem-loop, and unpaired bases). The stem had 2 bps and an UNCG tetraloop. **b**. Interaction between cPRT Stop (in cartoon) and bRT (surface with palm in green and thumb in red). **c**. Schematic of *c*PRT Stop duplex, in same format as Fig. 2c. Solid thick black lines indicate tertiary A-minor motif contacts. ^*avd*^A369 is a guanosine in RNA^Δ98^, introduced by *in vitro* transcription, and forms a wobble base pair with ^Sp^A70. The figure depicts the wild-type adenosine in gray. **d and e**. Tertiary A-minor motif interactions ofSPA80 with ^Sp^C69:^*avd*^G70 (d) and ^Sp^A81 with ^Sp^C68:^*avd*^G371 (e). ^SP^A80 and ^SP^A81 are also contacted by bRT. Black dotted lines are hydrogen bonds.

In the Sp strand, only two disordered nucleotides (^Sp^57 and 58) separated the *avd*:Sp duplex from the *TR*:Sp template:primer duplex. In the other strand, ∼30 disordered nts separated the ends visible in RNA^Δ98^ (between *avd*^381^ and *TR*^117^); in intact *TR*, this span would be ∼130 nts. These ∼30 nts were likely looped out between the *avd*:Sp and template:primer duplexes. In this architecture, the bulk of *TR* would extend towards the wide end of the Avd barrel (Fig. S9d), to which Fae was fused, explaining why the RNP containing Fae reverse transcribed RNA^Δ98^ with its truncated *TR* but not DGR RNA with its intact *TR* (Fig. S1b).

Significantly, *c*PRT of DGR RNA terminates within ^*avd*^383-387 (*9*), just before the *avd*:Sp duplex, suggesting that this duplex along with its associated elements (i.e., stem-loop and unpaired nucleotides) create a physical barrier to polymerization. We therefore termed these elements the ‘*c*PRT Stop’. Notably, deletion of the *avd* strand (Δ^*avd*^368-377) results not in longer cDNAs but instead no reverse transcription at all (*9*). This is not due to loss of RNA binding, as a DGR RNA lacking the *avd* segment still bound bRT and Avd (Fig. S10). This result suggested that the *c*PRT Stop not only terminates polymerization but is also required for *c*PRT. We asked whether base pairing in the *avd*:Sp duplex was essential for *<u>c</u>*<u>PRT</u>. Substitutions that disrupted 12 bps between *avd* and Sp resulted in loss of *c*PRT, but surprisingly reversion through complementary substitutions did not restore *c*PRT (Fig. S11). Disruption of just the six Watson-Crick base pairs had a similar result (Fig. S11). These findings suggested that the function of the *avd*:Sp duplex in *c*PRT is conformationally sensitive, possibly due to A-minor motif interactions, which were affected by both sets of mutations. In support of this conclusion, removal of the short stem-loop (Δ^Sp^71-78) preceding the A-minor motif interactions also led to loss of *c*PRT (Figs. 3a and S12). Together, these results suggested a structurally dynamic link between the *c*PRT Stop element and template:primer duplex, with proper alignment between the two required for *c*PRT.

### Thumb Ring, bRT-binding, and Avd-binding RNA

The *c*PRT Stop element continued in the Sp strand into two further RNA elements, both of which contacted bRT and were in density of excellent local resolution (Figs. 4 and S13). The first element (^Sp^82-94) wrapped around the base of the bRT thumb and was therefore called the ‘Thumb Ring’, and was notable in being the longest stretch of single-stranded RNA in the RNP. Eight of 12 unpaired bases in the Thumb Ring were contacted by bRT (Fig. 4b). Of these, ^Sp^C92 and U94 were the most significant for *c*PRT (Figs. 4c and S13). Substitutions at other bases in the Thumb Ring likewise reduced *c*PRT.

**Fig. 4.**
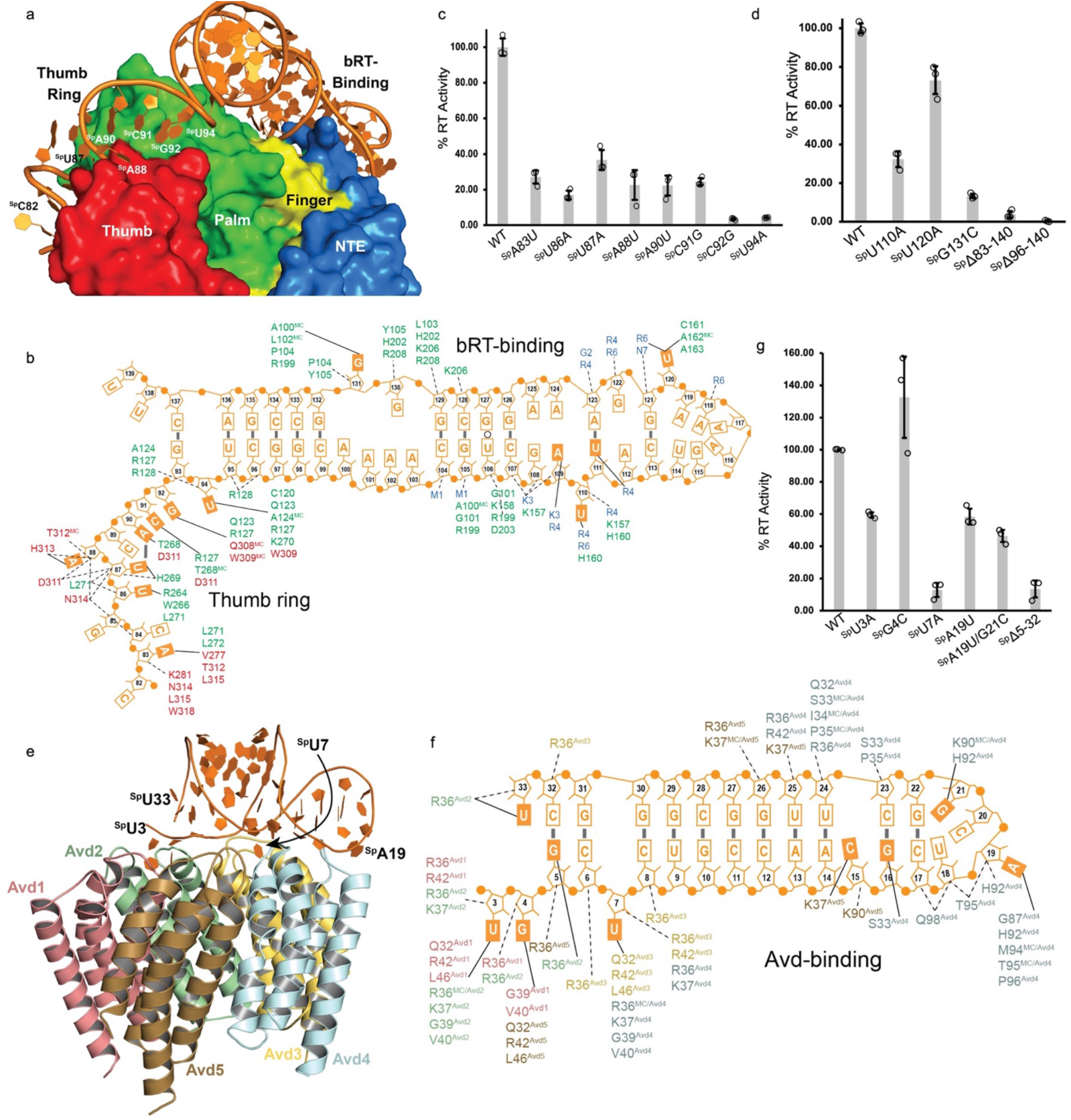
Thumb Ring, bRT-binding, and Avd-binding RNA. **a**. Interaction of Thumb Ring and bRT-binding RNA with the surface of bRT. **b**. Schematic of Thumb Ring and bRT-binding RNA, in same format as Fig. 2c. **c and d**. *c*PRT activity ofDGR RNA mutated in the Thumb Ring (c) or bRT-binding RNA element (d) relative to wild-type DGR RNA. Data here and below are from experiments that were repeated three independent times, and means and standard deviations shown. **e**. Interaction between Avd-binding RNA and pentameric Avd. **f**. Schematic of Avdbinding RNA, in same format as Fig. 2c. **g**. *c*PRT activity ofDGR RNA mutated in the Avd-binding RNA element relative to wild-type DGR RNA.

Sp continued from the Thumb Ring into a bRT-binding element composed of a long stem (^Sp^95-113 and 121-139) interrupted by two interior loops (^Sp^100-103:130-131 and ^Sp^108-112:122-125), and capped by a heptaloop (^Sp^114-120) (Figs. 4 and S12), the most common type of loop after tetraloops (*25*). The interior loops mostly had non-standard base-base interactions. Bases flipped-out from the stem (^Sp^U110 and G131) and heptaloop (^Sp^U120) were contacted by bRT palm and NTE. Substitutions at these bases decreased *c*PRT, with the greatest effect at ^Sp^G131 (Fig. 4d and S13). Deletion of the Thumb Ring along with the bRT-binding stem-loop (Δ^Sp^83-140) or the bRT-binding stem-loop alone (Δ^Sp^96-140) eliminated *c*PRT almost entirely. The latter also resulted in loss of RNA binding to bRT while binding to Avd was maintained (Fig. S10). bRT associates with Avd in the absence of RNA (*9, 12*), but this last result shows that this interaction is strengthened by DGR RNA.

The final RNA element consisted of the 5’ end of Sp (^Sp^3-33), which formed a 11-bp stem capped by a UNCG tetraloop (Figs. 4e, f and S14). This stem-loop was contacted by all five Avd protomers, consistent with RNase protection experiments (*9*). Bases flipped-out from the stem-loop were inserted into crevices between protomers (Fig. S14). The same arsenal of Avd amino acids but from different protomers contacted the flipped-out bases ^Sp^U3, G4, and U7. ^Sp^U7 made the most crucial contacts as its substitution reduced *c*PRT to the same level as deletion of the entire Avd-binding segment (Δ^Sp^5-32) (Figs. 4g and S11). Substitution of other flipped-out bases decreased *c*PRT less dramatically.

### RNA Conformational Variability

Several RNA elements — the template:primer duplex, *c*PRT Stop, and Thumb Ring — were conformationally variable, whereas others — bRT- and Avd-binding stem-loops — were stable across several alternative RNP conformations. These alternative conformations were evident in multiple samples and independent of incubation with dNTPs or analogs.

An RNP conformation in which the thumb was collapsed into the active site, called ‘Resting’, was visualized in three independent samples (Figs. 5a and S15, Table S1, Complexes 4-6), including one in which no dNTP or analog was added. The thumb was highly mobile and occupied both N- and T-sites, having displaced the disordered fingertips loop. The template:primer duplex skirted the surface of bRT rather than entering the active site. The bRT- and Avd-binding stem-loops were in the same positions as in the Active conformation, but density was lacking for the *c*PRT Stop and Thumb Ring in two of the samples, although in the third (Complex 5) these could be modeled into density of low local resolution (4-5 Å), which revealed that these RNA structures were disengaged from bRT.

**Fig. 5.**
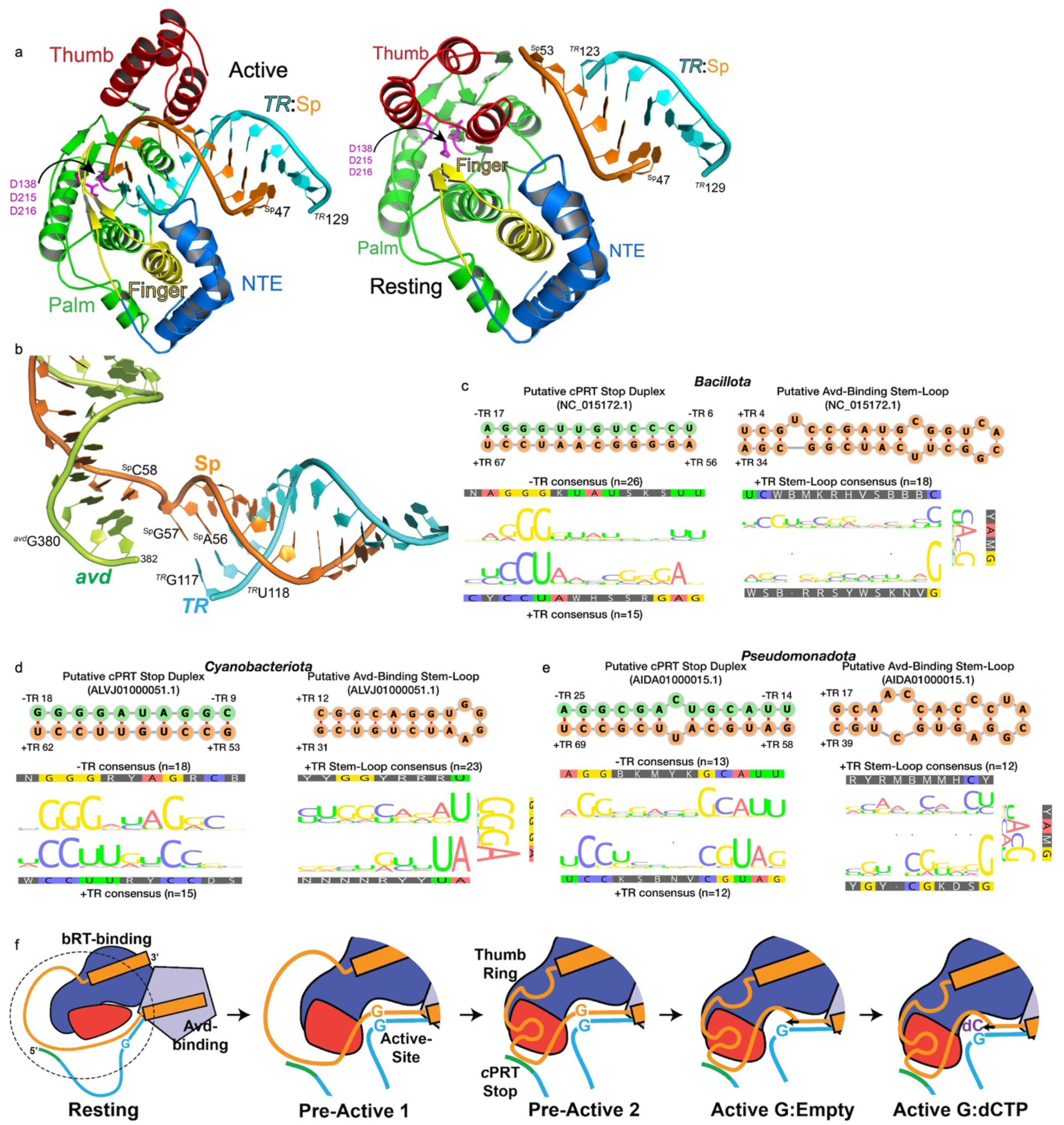
RNA Conformational Variability. **a**. (Left) Active and (right) Resting conformations, with bRT and *TR*:Sp template:primer duplex shown (coloring as in Fig. 2a). The locations of catalytic D 13 8, D215, and D216 identify the active site. The thumb subdomain was translated ∼ 19 Å and rotated ∼117° in the Resting conformation relative to the Active conformation. **b**. *TR* and Sp RNA in the Pre-Active I conformation. **c-e**. Representative RNA structure predictions from *Bacillota* (c), *Cyanobacteriota* (d), and *Pseudomonadota* (e) DGRs. Putative *c*PRT Stop duplexes and Avd-binding stem-loops are represented by example sequences (sequences upstream and downstream of *TR* are green and orange, respectively), with corresponding sequence alignment consensus profiles. The number of genomes in each alignment is indicated. **f**. Schematic of bRT-Avd:RNA conformational states, shown as a pathway from Resting to Active G:dCTP. The circled area in the Resting conformation is magnified in the remaining panels.

A second alternative RNP conformation, in which the *TR*:Sp duplex was in the active site but ^Sp^G57 occupied the N-site and ^Sp^C58 joined the *c*PRT Stop duplex, was visualized in three samples (Figs. 5b and S16, Table S1, Complexes 7-9), again including one in which no dNTP or analog was added. ^Sp^G57 and ^Sp^C58 were disordered in the Active conformation but here were seen to directly connect the *c*PRT stop and template:primer duplexes. ^Sp^G57, which is required for *c*PRT (*9*), formed a guanine-guanine pair with ^*TR*^G117 through their Watson-Crick faces.

While such interactions are rare, a similar pairing is observed in *E. coli* 16S rRNA (*26*). The Avd- and bRT-binding elements were in the same positions as in the Active conformation, but the stem-loop of the *c*PRT Stop was disengaged from bRT and the Thumb Ring was too flexible to be modeled. This conformation was called ‘Pre-Active 1’ to distinguish it from a variant in which ^Sp^G57 still occupied the N-site but the *c*PRT Stop stem-loop and Thumb Ring were positioned as in the Active conformation and engaged with bRT; this latter variant was called ‘Pre-Active 2’ (Fig. S17, Table S1, Complex 9). The Pre-Active 2 conformation appeared to be transitional between the Pre-Active 1 and Active conformations, as the density for ^Sp^G57 in the T-site was weaker in Pre-Active 2 than in Pre-Active 1 and ^Sp^C58 was too flexible to be modeled, consistent with ^Sp^G57 exiting the N-site and making it available for an incoming dNTP.

### Conservation of RNA elements

DGRs from microbial genomes spanning *Pseudomonadota, Bacillota, Cyanobacteriota, Nanoarchaeota*, and members of the Candidate Phylum Radiation (CPR) among others had RNA structural elements similar to ones observed in the BΦ DGR. Of 278 DGRs that have Avd homologs (*13*), 207 had putative Avd-binding stem-loops located in sequences just downstream of *TR* (Figs. 5c-e and Fig. S18, and Table S3). The stems most commonly had 9 bps, contained potentially flipped-out bases, and were capped by tetraloops. GNRA tetraloops were prevalent among *Cyanobacteriota* DGRs and UNCG tetraloops occurred frequently in *Pseudomonadota* DGRs. The same 207 DGRs had putative *c*PRT Stop duplexes, involving pairing of distal sequences (upstream and downstream of *TR*). The duplexes had an average of 13 bps, with sequences especially conserved in *Cyanobacteriota* DGRs. The *c*PRT Stop element is complex, involving not only a duplex but also a stem-loop followed by adenosines in the sequence downstream of *TR*. Remarkably, 47 DGRs had predicted stem-loops adjacent to the duplex, with all but two having multiple adenosines following, suggesting the presence of A-minor motif interactions. These results strongly suggested that the mechanism of DGR reverse transcription was conserved across distant taxa.

## Discussion

We sought to understand how the DGR RNA with its essential RNA segments surrounding *TR* combines with bRT and Avd to initiate reverse transcription. Most RTs require a primer for cDNA synthesis, and the source of that primer varies widely. For example, group II intron RTs and non-LTR retrotransposons use nicked target DNA as a primer (*21, 27-30*), HIV-1 RT uses a host tRNA (*31, 32*), and telomerase uses an associated RNA (*33*). These primers along with the corresponding templates are recognized directly through sequence, structure, or both.

Our work revealed an unprecedented mechanism for the BΦ DGR, in which the primer and template were not recognized directly, but rather placed precisely in the bRT active site through extensive, intricate, and intimate interaction of RNA structural elements surrounding the template.

Crucial structures and functions were distributed across the entire length of Sp RNA. At its 5’ and 3’ ends, Sp formed stem-loops that acted as grapples for Avd and bRT, respectively, binding to these proteins through flipped-out bases. The grapples were positioned similarly across multiple RNP conformations. These stable grapples provide a ready explanation for the processivity endowed by the DGR RNA, enabling *TR* to maintain association with bRT-Avd through multiple polymerization cycles. Notably, bRT and other DGR RTs lack the processivity-promoting α-loop found in group II intron RTs (Fig. S18) (*34*). In between the ends, the DGR RNA formed the template:primer duplex, *c*PRT Stop, and Thumb Ring. The latter two elements were especially mobile across multiple RNA conformations, and in the Active conformation interacted extensively with bRT, appearing to provide a conformational guide for the placement of the template:primer duplex in the bRT active site.

The multiple RNP conformations observed lay along a plausible pathway from initial association of the DGR RNA with bRT-Avd to initiation of reverse transcription. We surmise that the Resting conformation corresponds to initial association, with the template:primer duplex on the surface of bRT, blocked from entering the active site by the collapsed thumb subdomain, and the *c*PRT Stop and Thumb Ring disorganized and disengaged from bRT. A similar conformation with the thumb collapsed into the active site in the absence of a template:primer duplex occurs in HIV-1 RT (*35*). The Resting conformation likely proceeds to the Pre-Active 1 conformation, with the thumb receding and the template:primer duplex entering the active site, but with ^Sp^57G in the N-site. We had previously found that ^Sp^G57 was required for *c*PRT (*9*), but the basis for this was unexplained. Our current results suggest that guanosine at ^Sp^57 is necessary because it stabilizes the N-site prior to complete organization of Sp RNA into the Active conformation. As noted above, the Pre-Active 2 conformation likely represents a transition to the Active conformation, with the RNA completely organized and ^Sp^57G at the N-site but with greater apparent mobility. Displacement of ^Sp^57G from the N-site would result in the Active N-empty conformation, and finally, binding of an incoming dNTP would stabilize the Active conformation, enabling chemistry to occur. An N-empty conformation with adenosine at the T-site was nearly indistinguishable with one with guanosine. No contact by bRT or Avd was made to the substituent at the C6 position of A or G, which is a crucial determinant of A-mutagenesis (*14*). These observations suggest that A-mutagenesis is unlikely a result of specific protein contacts or major structural rearrangements, and more likely attributable to dynamics.

In summary, we discovered that reverse transcription in the BΦ DGR depends on an abundance of essential interactions between RNA structural elements and bRT-Avd, resulting in the precise positioning of a template:primer duplex for *cis*-priming and strictly restricting A-mutagenesis to DGR variable protein. Strikingly, one of the more complex RNA structural elements, the *c*PRT Stop, appeared to be present in DGRs from distant taxa, suggesting an extraordinary conservation of the mechanism of reverse transcription.

## Materials and Methods

### Protein expression and purification, RNA synthesis, and reverse transcription

The coding sequence for the Avd fusion partner *Methylobacterium extorquens* formaldehyde activating enzyme (Fae, PDB 1Y60, 170 amino acids) was synthesized with codons optimized for *E. coli* expression (Genewiz). The coding sequence, starting from the codon for Fae amino acid 4, was inserted after the Avd codon for amino acid 124. bRT and Avd or Avd-fusion protein were co-expressed in *E. coli* BL21 Gold (DE3) and purified as described (*9*). RNA was produced through *in vitro* transcription using T7 RNA polymerase and gel-purified, and reverse transcription reactions with bRT and Avd or Avd-fusion protein were carried out, as described (*9*). RNA and cDNA were examined by denaturing gel electrophoresis, as described (*9*), and in some cases, cDNA was subjected to short-read next-generation sequencing (Amplicon EZ, Genewiz), as described (*14*).

### Cryo-EM sample preparation and data collection

For assembly of complexes, 150 µL of RNA^Δ98^ (14 µM) was incubated in 1.2 mL of 75 mM KCl, 3 mM MgCl_2_, 10 mM DTT, 50 mM HEPES, pH 7.5, and 10 % glycerol at 37 °C for 15 min. bRT-Avd-Fae (150 µL of 14 µM) was added and the mixture was further incubated at 37 °C for 2 h. The concentration of the complex was determined using A_280_ measured with a NanoDrop (Thermo Scientific) and a calculated molar extinction coefficient of 206,960 M^-1^cm-^1^..The sample was passed through a 0.22 µm filter and concentrated to 100 µL using a 30-kDa molecular weight cutoff membrane. The complex was resolved by gel filtration at 4 °C with a Superdex 200 10/300 column (GE Healthcare) equilibrated with 75 mM KCl, 30 mM (NH_4_)_2_SO_4_, 50 mM Tris, pH 7.5, 3 mM MgCl_2_, and 2 mM DTT. Eluted fractions were analyzed by SDS-PAGE to visualize proteins, and denaturing gel electrophoresis (8 M urea containing 8 % polyacrylamide gels in 1× TBE buffer) to visualize RNA. Fractions containing bRT-Avd-Fae/RNA^Δ98^ complexes were concentrated to ∼2 µM using a 30-kDa molecular weight cutoff membrane. The concentration of bRT-Avd-Fae/RNA^Δ98^ was determined by measuring absorbance at A_260_ with a NanoDrop (Thermo Scientific), assuming a 1:5:1 bRT:Avd-Fae:RNA^Δ98^ stoichiometry. The concentration of the complex was adjusted to 1 μM and left either unincubated or incubated with the following dNTP analogs or dNTPs at 37 °C for 2 h. For complexes containing wild-type RNA^Δ98^, 100 µM of non-hydrolyzable dCTP (2’-deoxycytidine-5’-[(α,β)-methyleno]triphosphate, Jena Bioscience) was used. For complexes containing *TR* G117A RNA^Δ98^, 2 mM non-hydrolyzable dCTP (as above) or 2 mM non-hydrolyzable dTTP (2’-deoxythymidine-5’-[(α,β)-methyleno]triphosphate, Jena Bioscience) was used. For complexes containing *TR* U114A RNA^Δ98^, 2 mM each of dCTP, dATP, and ddGTP was used. Complexes (4 μL) were then applied manually 2-3x in quick succession to Quantifoil™ R 1.2/1.3 Au grids, which had been freshly plasma cleaned in Gatan Solarus II (under Ar/O_2_ mixture) for 10 sec.

After application, grids were blotted with varying force and times using a Vitrobot Mark IV (Thermo Fisher) at 4 °C and 100% humidity, and then immediately plunge-frozen in liquid ethane.

Cryo-EM data were collected on either a 300 keV Titan Krios cryo-transmission electron microscope at the Pacific Northwest Center for Cryo-EM (PNCC) equipped with GIF quantum energy filters attached to either a K3 electron counting direct detector (Gatan) using Serial EM software, or a 300 keV Titan Krios cryo-transmission electron microscopes (FEI Company) at the University of California San Diego Cryo-EM facility equipped with GIF quantum energy filters attached to a K2 electron counting direct detector (Gatan) using EPU (Thermo Fisher).

Images were acquired at a magnification of ∼130,000, corresponding to a calibrated raw pixel size of ∼0.5 Å, in EFTEM mode with an energy filter slit width of 10-20 eV. Movie image stacks with a total electron dose of ∼55 e^−^ /Å^2^ were saved in non-super resolution counting mode.

### Cryo-EM data processing and analysis

The collection and processing of raw data is outlined in detail in Fig. S2 and Table S1. Briefly, raw movies of each independent dataset were imported in cryoSPARC 3.1.1 (*36*), RELION 3.0, or both (*37*); motion-corrected using patch motion correction (multi) of cryoSPARC or MotionCor2; and CTF estimated using patch CTF estimation (multi) in cryoSPARC, or GCTF or CTFFIND4 in RELION. A list of micrographs for each dataset (removing poor quality micrographs using various criteria e.g., CTF fit of < 8 Å, astigmatism < 1200, motion distribution < 60, and motion curvature < 40) was prepared. Data were collected at a relative ice thickness of ∼1-1.2 μm, as thin ice led to particle denaturation whereas thick ice led to high background noise. Particles were picked from curated micrographs using various methods and iterative strategies outlined in Fig. S2a. Duplicate particles were removed in cryoSPARC, and binned or un-binned particles were extracted from micrographs depending on the purpose and stage of analysis. Multiple rounds of 2D classification were performed to eliminate poor quality particles that could not be classified (Fig. S3). Selected particles from 2D classes were used for initial 3D classification (with different number of classes) using *ab initio* reconstruction (Fig. S4). Particles belonging to poorly resolved *ab initio* 3D class averages were discarded. Particles from 3D classes which indicated the presence of all domains were used for heterogeneous classification with various plausible models. Good classes from heterogeneous refinements were fed back into refined template picking using programs outlined in Fig. S2.

Iteration of the above processes provided a refined set of particles, which were classified into probable 3D classes (judged from shape profiles of 3D models of heterogenous classification), and eventually particles from final 3D class sets were extracted without binning with a 320-pixel box size for consecutive refinement using homogeneous, non-uniform, and local refinement algorithms in cryoSPARC with appropriate masks. Per-particle defocus refinement was also performed, although resulting in little to no discernible improvement in map quality. Trials of particle subtraction did not indicate any improvement of maps and thus this step was omitted from refinement. The total number of movies, number of particles used for final refinement of each class, orientation distribution, half-map-based correlation, global resolution estimate, and several other parameters are provided in Table S1. Density sharpened versions of the maps were used for model building.

### Model building and refinement

The model building pipeline is depicted in detail in Fig. S2b. In brief, an initial model of bRT was built manually using PDB templates of poliovirus RNA-dependent RNA polymerase (3OL9) and group II intron RT GsI-IIC (6AR1), and the initial model of Avd was based on the structure of the free form of the pentamer (PDB 4DWL). For modeling RNA, the secondary structure for DGR RNA was first predicted through iterative rationalization using RNAstructure (*38*), incorporating refined SHAPE-MaP data and biochemical results, and finalized using initial map features (e.g., base pairing in the *TR*:Sp template:primer duplex) (*9, 15*). The dot bracket files defining plausible RNA stem regions were used with simRNA (*39*) and RNAComposer (*40*) to obtain initial 3D templates of RNA stems, which were docked in the sharpened maps. Rigid body fitted models of bRT subdomains, Avd protomers, RNA stems and the connecting single stranded RNA regions were iteratively (re)built in real space in Coot (*41*). In addition, a pentameric model of Fae (PDB 1Y60) was placed into the diffuse density contiguous to the density for the Avd pentamer. The models of complexes composed of bRT, Avd, and RNA^Δ98^ were finally refined in CCPEM (*42*). Model geometry was validated using MolProbity package within the Phenix suite (*43*). This final model displays good stereochemistry (Table S1), except for regions of high flexibility. RNA structure was analyzed using RNAPdbee (*44*). Q-scores (*45*) were computed using Chimera (*46*). Derived B-factors were calculated as 150 • (1 – Q) (*18*), with the scale factor of 150 empirically identified to provide the highest map correlation coefficient, as calculated with Phenix (*43*) (Table S2). Figures of molecular models were made with PyMOL (The PyMOL Molecular Graphics System, Schrödinger, LLC) and ChimeraX (*47*).

### SHAPE-MaP

SHAPE-MaP experiments were carried out using 10 mM N-methylisatoic anhydride (NMIA, Millipore-Sigma). Thirty-five µL of DGR RNA (500 ng/µL) in water was denatured at 95 °C for 3 min and then immediately placed on ice for 5 min. The RNA was refolded by incubation at 37 °C for 20 min in 110 mM KCl, 70 mM HEPES, pH 7.5, 4 mM MgCl_2_ in a total volume of 270 µL. Five µL of 300 mM (NH_4_)_2_SO_4_, 20 mM HEPES, pH 7.5, 5 mM MgCl_2_, 2 mM DTT, and 10% glycerol containing bRT-Avd (2 mg/mL) was added to 40 µL of refolded RNA, whereupon the sample was incubated at 37 °C for 1 h. Five µL NMIA (100 mM in DMSO) was then added to the samples, which were further incubated at 37 °C for 1 h. After this time, NMIA was quenched by the addition of DTT to a final concentration of 50 mM. A negative control for the reaction was carried out similarly, but with 5 µL of DMSO instead of NMIA. Reactions were then incubated with 2 µL of proteinase K (Ambion, 0.4 mg/ml) at 50 °C for 30 min, and proteinase K was then quenched with a final concentration of 1 mM PMSF. The resulting samples were purified with a desalting column (G-25).

A denaturing control was carried out by incubating 2 µL of RNA (500 ng/µL), 10 µL formamide, 2 µL of 10x denaturing buffer (500 mM HEPES, pH 8.0, and 40 mM EDTA) in a total volume of 18 µL. The sample was incubated at 95 °C for 1 min, and 2 µL of NMIA (100 mM) was added and further incubated at 95 °C for 1 min. The sample was placed on ice for 5 min, and then purified with a desalting column (G-25) after adjusting the volume to 50 µL.

Reverse transcription reactions were carried out by first annealing 8 µL of purified RNA from the previous step with 1 µL of reverse transcription primer P1 (2 µM, GACTGGAGTTC AGACGTGTGCTCTTCCGATCTGAAGTCGGCCCCGCCTTT) and 3 µL water at 65 °C for 5 min and cooling on ice for 5 min. To these annealed reactions, 8 µL of 2.5x MaP buffer (50 mM Tris, pH 8.0, 75 mM KCl, 6 mM MnCl2, 10 mM DTT, and 0.5 mM dNTPs) and 1 µL of ProtoScript^®^ II Reverse Transcriptase (NEB) was added. These reactions were incubated at 42 °C for 3 h and then at 70 °C for 15 min. These samples were then purified with a desalting column (G-25) after adjusting the volume to 50 µL.

The cDNAs resulting from the reverse transcription reactions were amplified in a 100 µL PCR reaction using 10 µL of the purified sample from the previous step, 50 µL of Q5^®^ High-Fidelity 2X Master Mix (NEB), 5 µL each of 10 µM primer P1 and P2 (ACACTCTTTCCCTACACGACGCTCTTCCGATCTAAGGGCA GGCTGGGAAAT). The PCR conditions used for amplification were: 98 °C for 30 s; 30 cycles of 98 °C for 10 s, 60 °C for 30 s, and 72 °C for 20s; followed by 72 °C for 2 min. The amplified DNA was gel purified and the samples were deep-sequenced in multiplexes using the Illumina MiSeq platform (2 × 250 bp, paired-end, Genewiz). Demultiplexed fastq files were processed using ShapeMapper 2.0 pipeline (*48*).

### EMSA and Mass Spectrometry

Reverse transcription was carried out at 37 °C for 20 min in 20 µL volume containing 1.4 µM RNA, varying concentrations of bRT-Avd (0-2.0 µM), and 20 units RNase inhibitor (NEB) in 75 mM KCl, 3 mM MgCl_2_, 10 mM DTT, 50 mM HEPES, pH 7.5, 10% glycerol. The reaction contents were centrifuged at 12,000 x *g* for 10 min, and 15 µL was loaded on a native 6% polyacrylamide gel and resolved for 90 min at 100 V. The gel was stained with GreenGlo™ Safe DNA Dye (Thomas Scientific) and imaged on a Bio-Rad Chemidoc™ system. Gel-shifted bands (Fig. S13, black arrow) were extracted from the gel, and subjected to mass spectrometry on an Orbitrap Fusion Lumos MS at the UCSD Biomolecular and Proteomics Mass Spectrometry facility. Mass spectrometry data were analyzed using Peaks Studio 10.6 software.

### Analysis of RNA elements

The subset of DGR sequences from Wu et al. (*13*) that contain an Avd-like gene along with five archaeal genomes were analyzed. For each sequence, 40-nt upstream of *TR* was extracted and concatenated to 100-nt downstream of *TR* (i.e. spanning most of the putative Sp). RNA fold predictions were carried out with MxFold2 tool v. 2.4.18 (*49*). The resulting structural predictions were inspected for putative *cPRT* Stop duplexes (complementarity between sequences up- and downstream of *TR*) and Avd-binding stem-loops (in sequences downstream of *TR*).

## Acknowledgments

Use of Pacific Northwest Center for Cryo-EM (PNCC) supported by NIH grant U24GM129547. We thank R. Baker and A. Leschziner for initial cryo-EM characterization of complexes, T. Baker for computational resources, and J. Meyers for data collection at PNCC.

## Author Contributions

Conceptualization: SH and PG

Investigation: SH, TB, JC, BGP, PG

Visualization: SH, TB, BGP, PG

Funding acquisition: PG, GG

Project Administration: PG

Supervision: PG

Writing – original draft: SH, TB, BGP, PG

Writing – review & editing: SH, TB, GG, BGP, PG

## Supplementary Information

Supplementary Figures S1-S19. Supplementary Tables S1-S3.

