## Supplementary Figures S1-S19 for "Structural Requirements for Reverse Transcription by a Diversity-generating Retroelement"

Figure S1

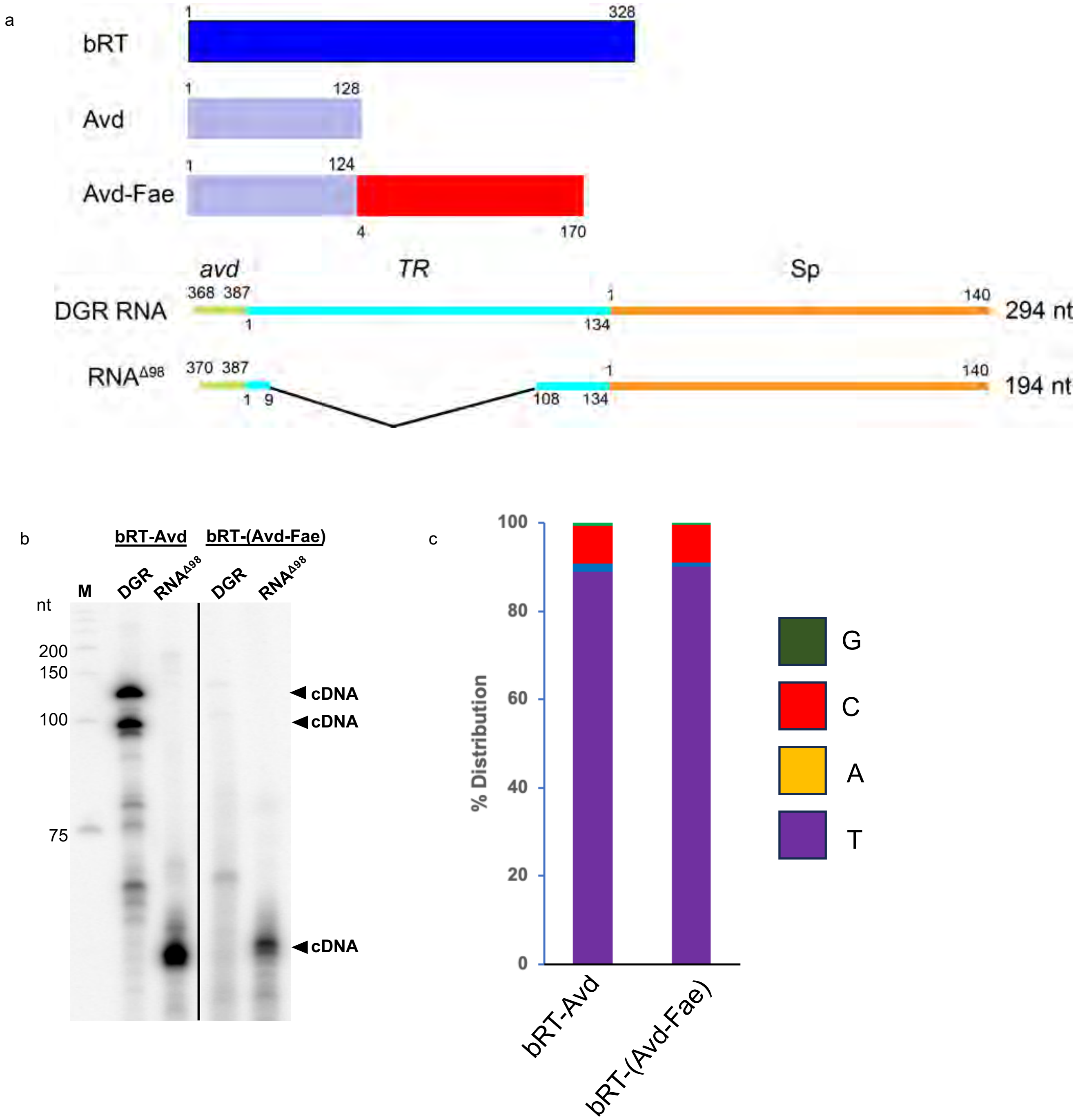


**Fig. S1. bRT-(Avd-Fae):RNA$^{\Delta98}$. a.** Schematic of bRT, Avd, Avd-Fae and DGR RNA and RNA$^{\Delta98}$. **b.** Synthesis of cDNA by bRT-Avd or bRT-(Avd-Fae) from DGR RNA or RNA$^{\Delta98}$. Reactions were incubated with dNTPs, including [α-$^{32}$P]dCTP, for 2 h at 37 °C. Products were treated with RNase and resolved on 8% denaturing polyacrylamide gel electrophoresis (PAGE). Lane M corresponds to radiolabeled, single-stranded DNA molecular mass markers. The panels are from the same gel in which irrelevant data were deleted (marked by line). **c**. Frequency distribution of deoxynucleotides incorporated (thymidine, purple) or misincorporated (adenosine, orange; cytosine, red; guanosine, green) by bRT-Avd (T 88.9%, A 1.8%, C 8.7%, G 0.6%) or bRT-Avd-Fae (T 90.0%, A 0.9%, C 8.7%, G 0.4%) at $^{TR}$G117A in RNA$^{\Delta98}$. $^{TR}$G117A has a lower misincorporation frequency than other adenosines in *TR* (1).

Figure S2

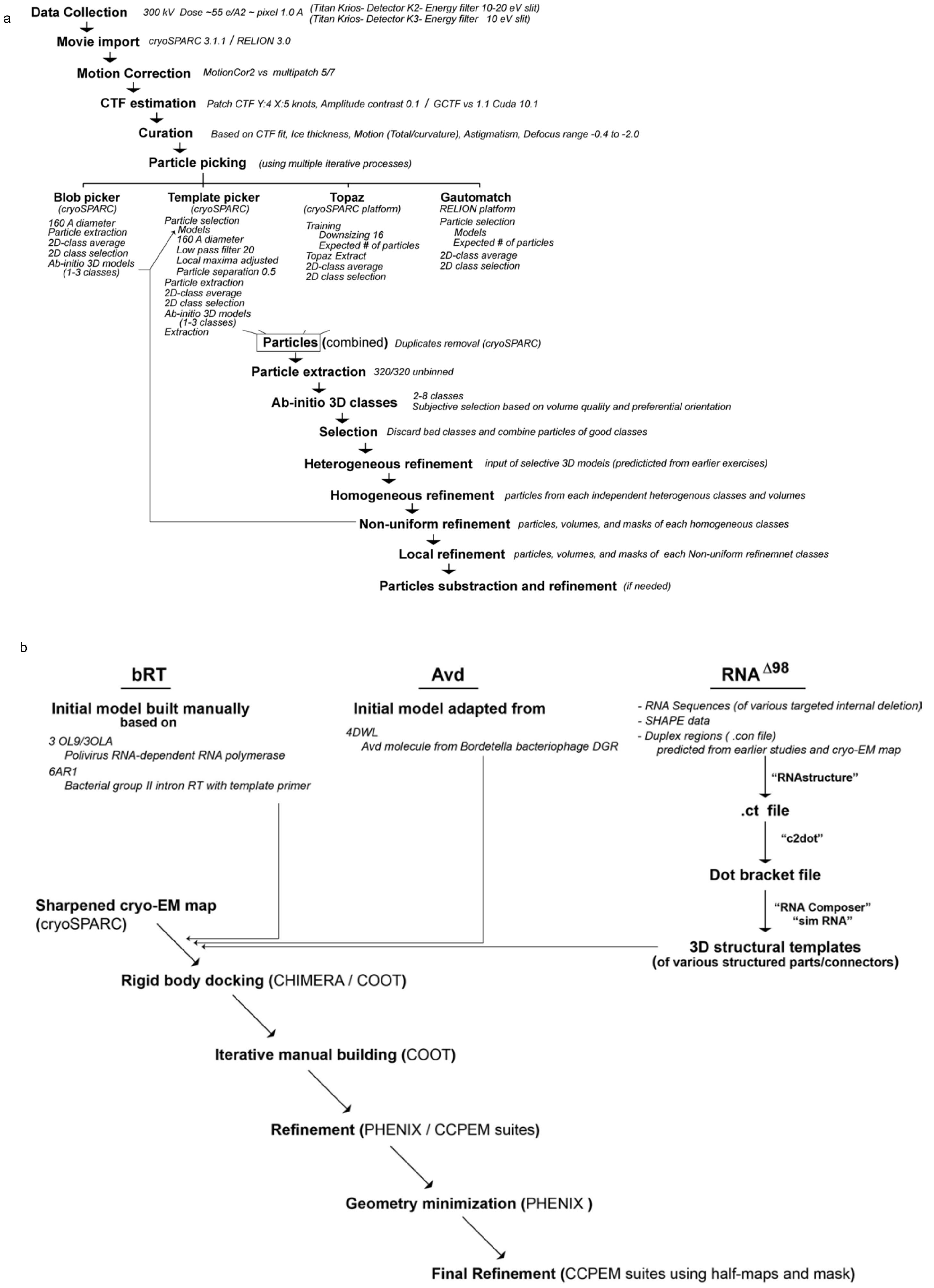
a
Data Collection
300 kV Dose ~55 e/A2 ~ pixel 1.0 A
(Titan Krios- Detector K2- Energy filter 10-20 eV slit)
(Titan Krios- Detector K3- Energy filter 10 eV slit)
Movie import
cryoSPARC 3.1.1 / RELION 3.0
Motion Correction
MotionCor2 vs multipatch 5/7
CTF estimation
Patch CTF Y:4 X:5 knots, Amplitude contrast 0.1 / GCTF vs 1.1 Cuda 10.1
Curation
Based on CTF fit, Ice thickness, Motion (Total/curvature), Astigmatism, Defocus range -0.4 to -2.0
Particle picking
(using multiple iterative processes)
Blob picker
(cryoSPARC)
160 A diameter
Particle extraction
2D-class average
2D class selection
Ab-initio 3D models
(1-3 classes)
Template picker
(cryoSPARC)
Particle selection
Models
160 A diameter
Low pass filter 20
Local maxima adjusted
Particle separation 0.5
Particle extraction
2D-class average
2D class selection
Ab-initio 3D models
(1-3 classes)
Extraction
Topaz
(cryoSPARC platform)
Training
Downsizing 16
Expected # of particles
Topaz Extract
2D-class average
2D class selection
Gautomatch
RELION platform
Particle selection
Models
Expected # of particles
2D-class average
2D class selection
Particles (combined)
Duplicates removal (cryoSPARC)
Particle extraction
320/320 unbinned
Ab-initio 3D classes
2-8 classes
Subjective selection based on volume quality and preferential orientation
Selection
Discard bad classes and combine particles of good classes
Heterogeneous refinement
input of selective 3D models (predicticted from earlier exercises)
Homogeneous refinement
particles from each independent heterogenous classes and volumes
Non-uniform refinement
particles, volumes, and masks of each homogeneous classes
Local refinement
particles, volumes, and masks of each Non-uniform refinemnet classes
Particles substraction and refinement
(if needed)
b
bRT
Initial model built manually
based on
3 OL9/3OLA
Polivirus RNA-dependent RNA polymerase
6AR1
Bacterial group II intron RT with template primer
Avd
Initial model adapted from
4DWL
Avd molecule from Bordetella bacteriophage DGR
RNA Δ98
- RNA Sequences (of various targeted internal deletion)
- SHAPE data
- Duplex regions ( .con file)
predicted from earlier studies and cryo-EM map
“RNAstructure”
.ct file
“c2dot”
Dot bracket file
“RNA Composer”
“sim RNA”
3D structural templates
(of various structured parts/connectors)
Sharpened cryo-EM map
(cryoSPARC)
Rigid body docking (CHIMERA / COOT)
Iterative manual building (COOT)
Refinement (PHENIX / CCPEM suites)
Geometry minimization (PHENIX )
Final Refinement (CCPEM suites using half-maps and mask)

**Fig. S2. Pipeline for cryo-EM data processing and map generation. a.** Flow-chart indicating the steps for collecting cryo-EM data, importing and processing movies, selecting particles belonging to unique 3D classes, and generating and refining maps. **b.** Schematic of the atomic model building and refinement processes for bRT, Avd, and RNA$^{\Delta 98}$.

Figure S3

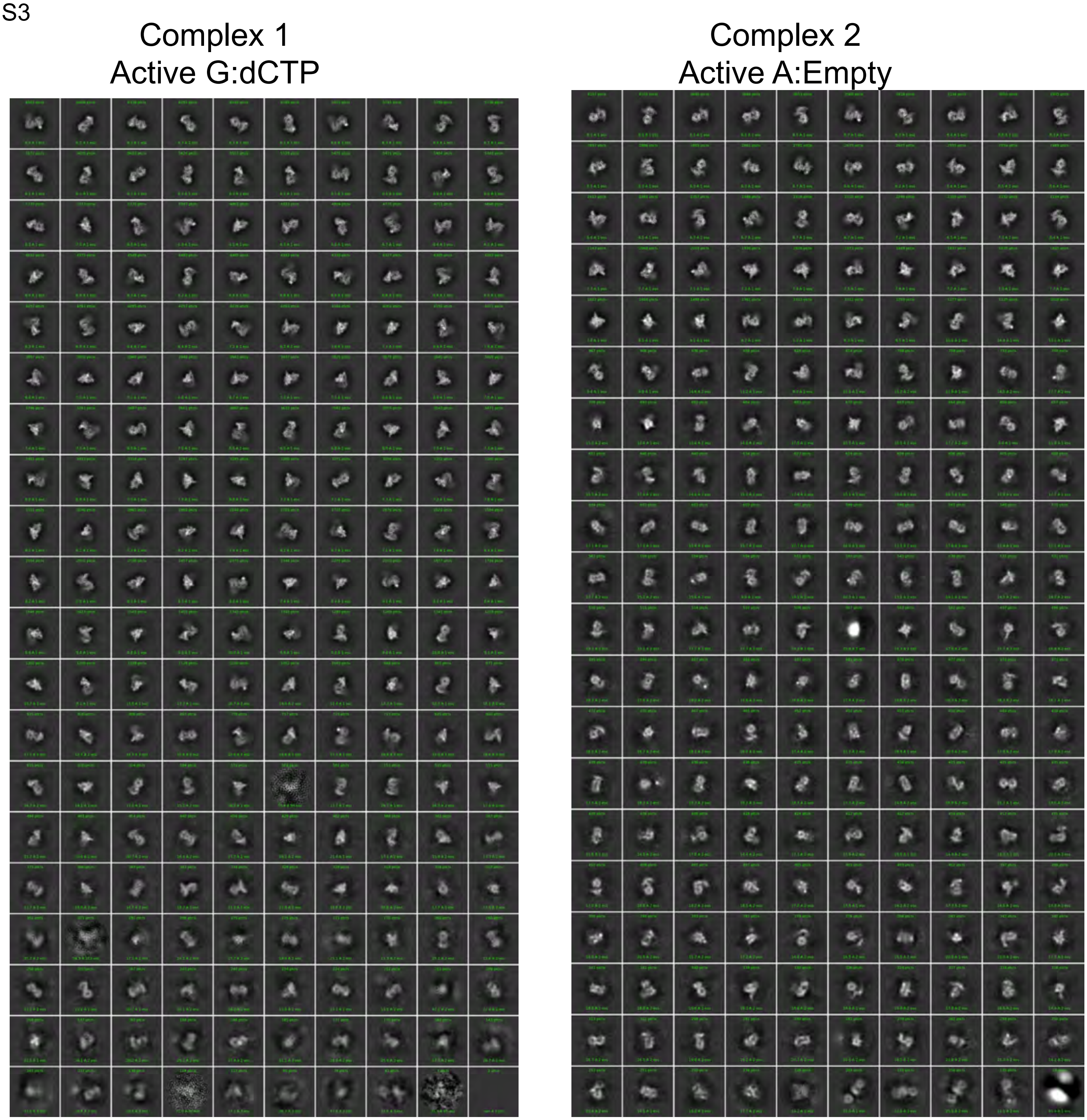


**Fig. S3. 2D Classifications.** Accepted 2D classification from cryoSPARC for 9 complexes (numbered as in Table S1) corresponding to 3D class separations and refinement processes.

Figure S3, continued

Complex 3
Active G:Empty

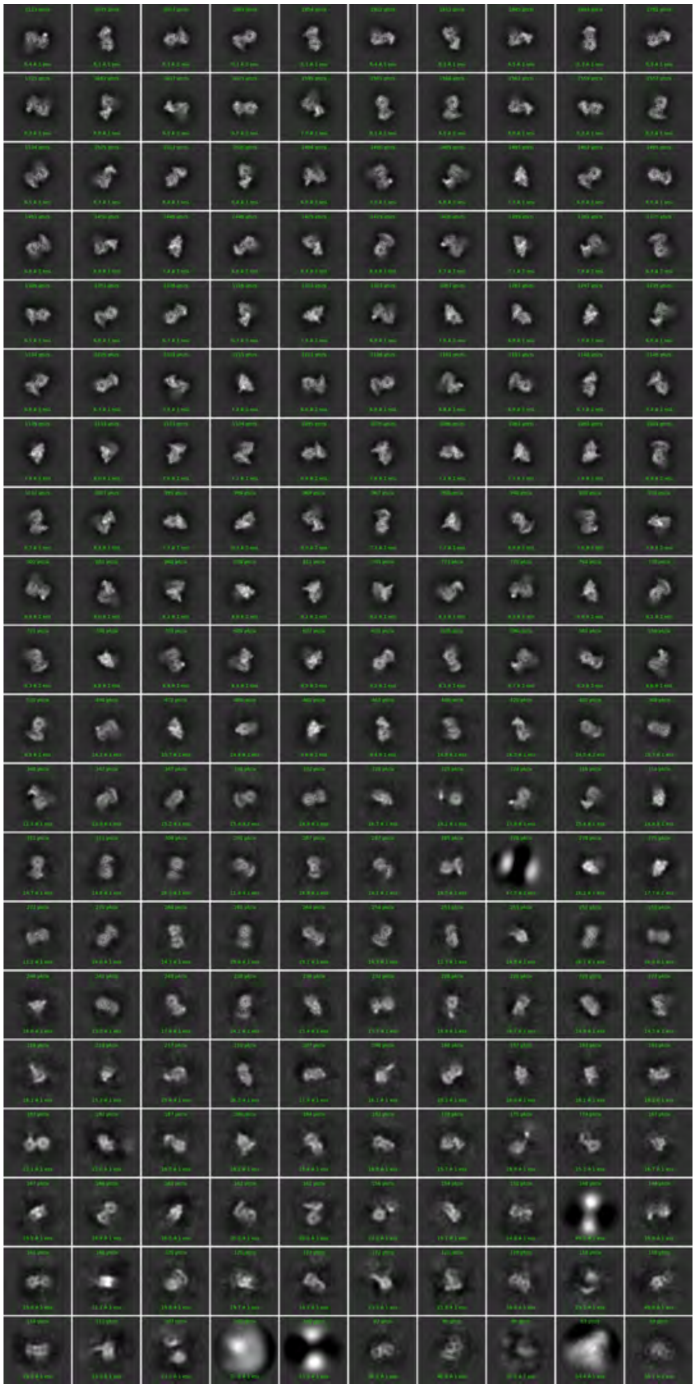

Complex 4
Resting

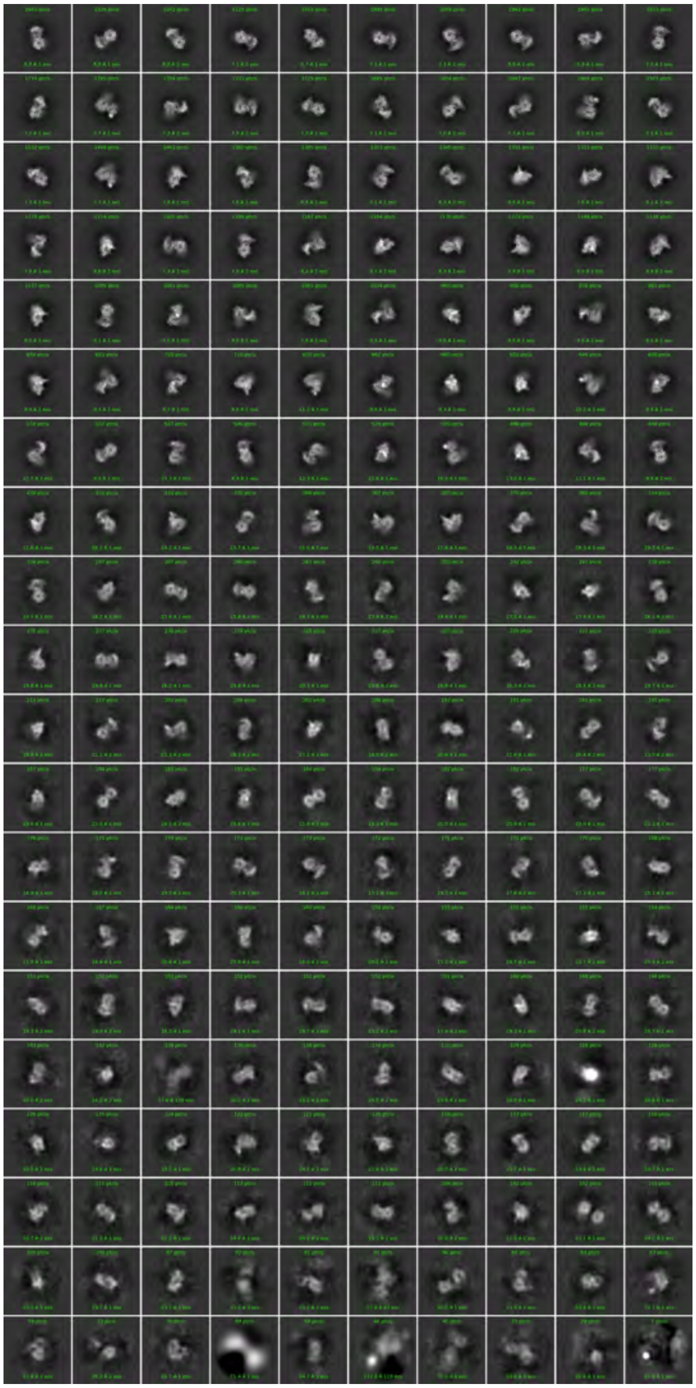

Figure S3, continued

Complex 5
Resting

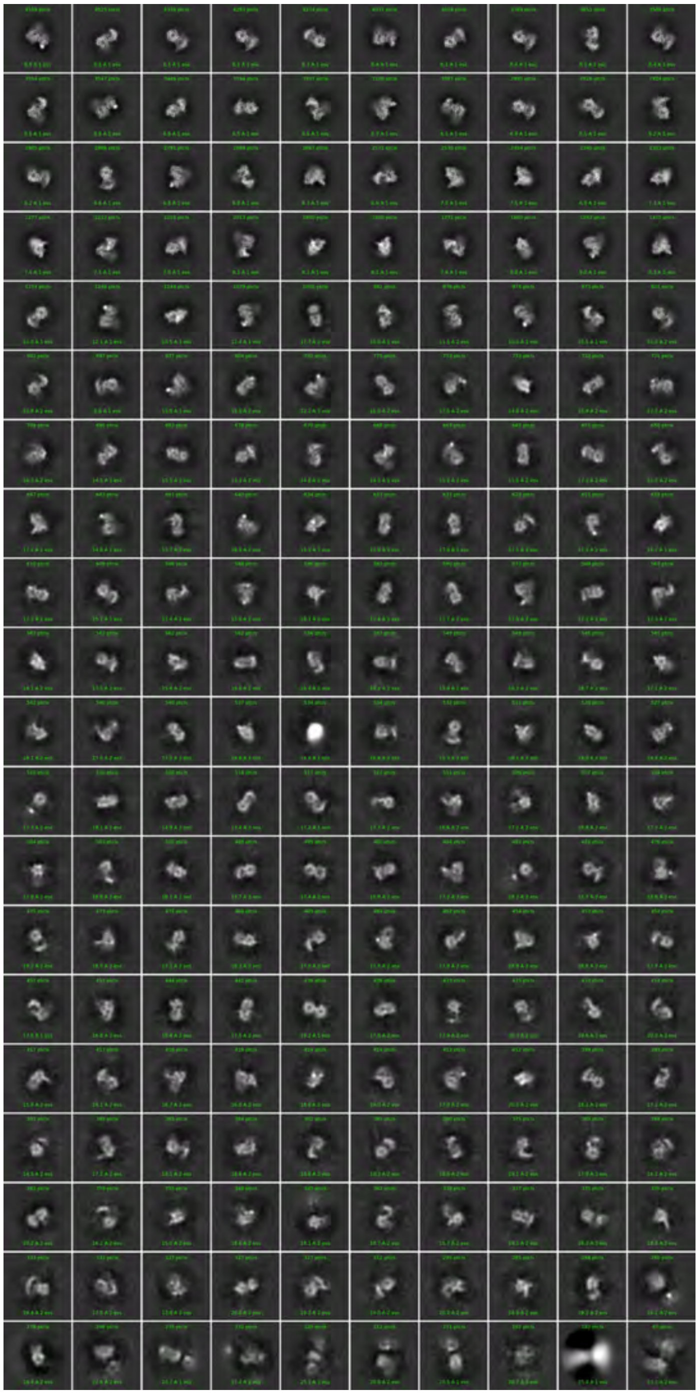

Complex 6
Resting

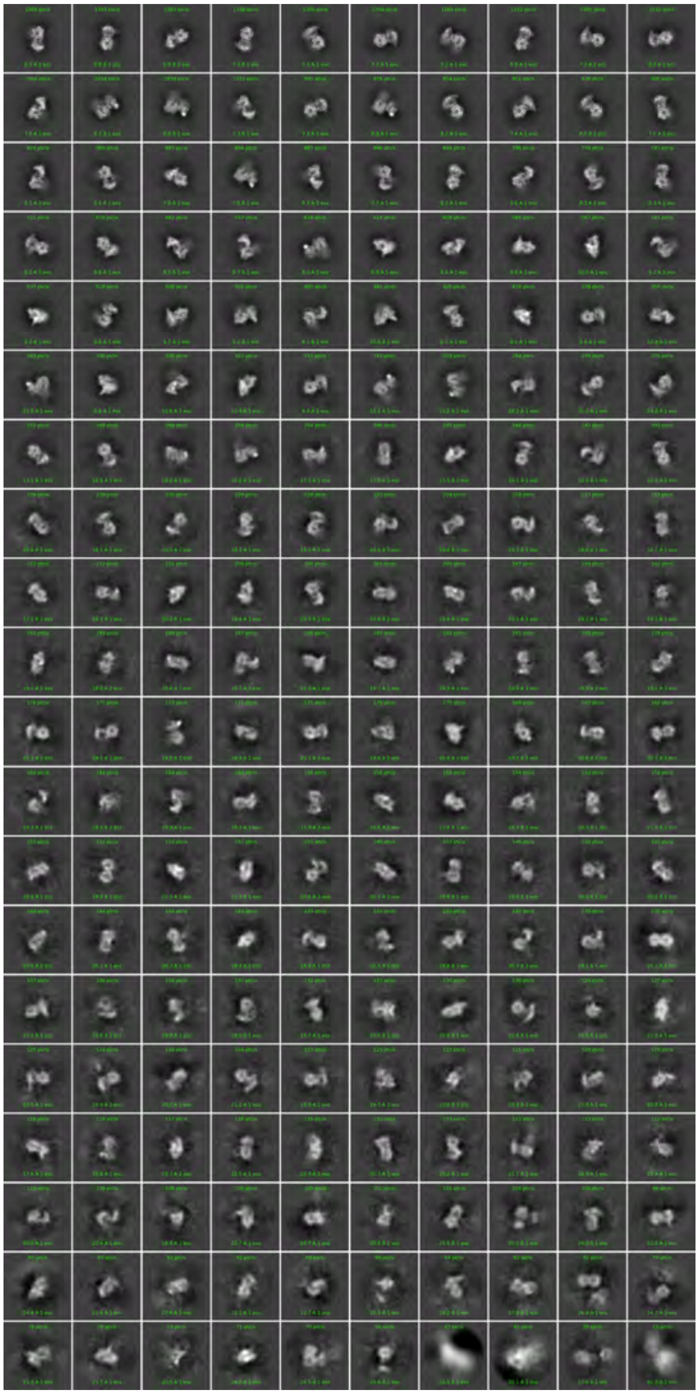

Figure S3, continued

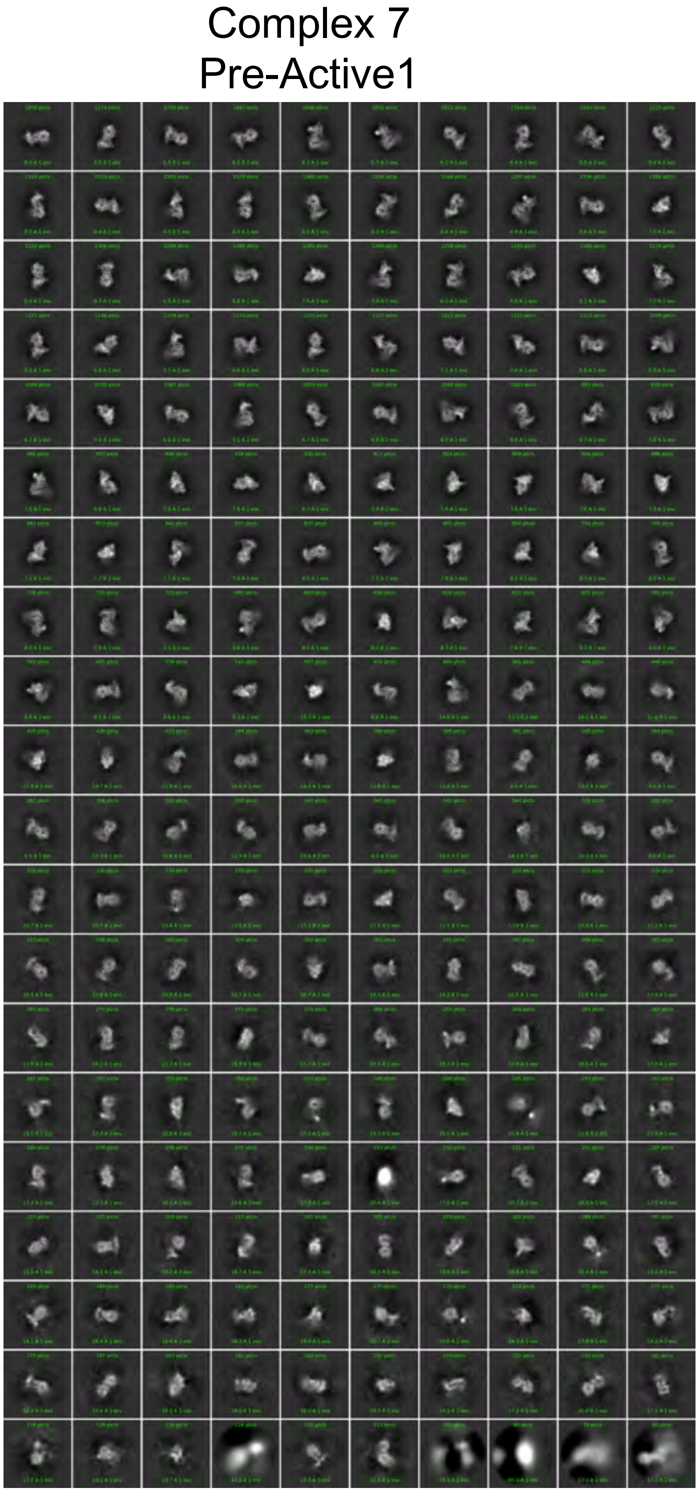

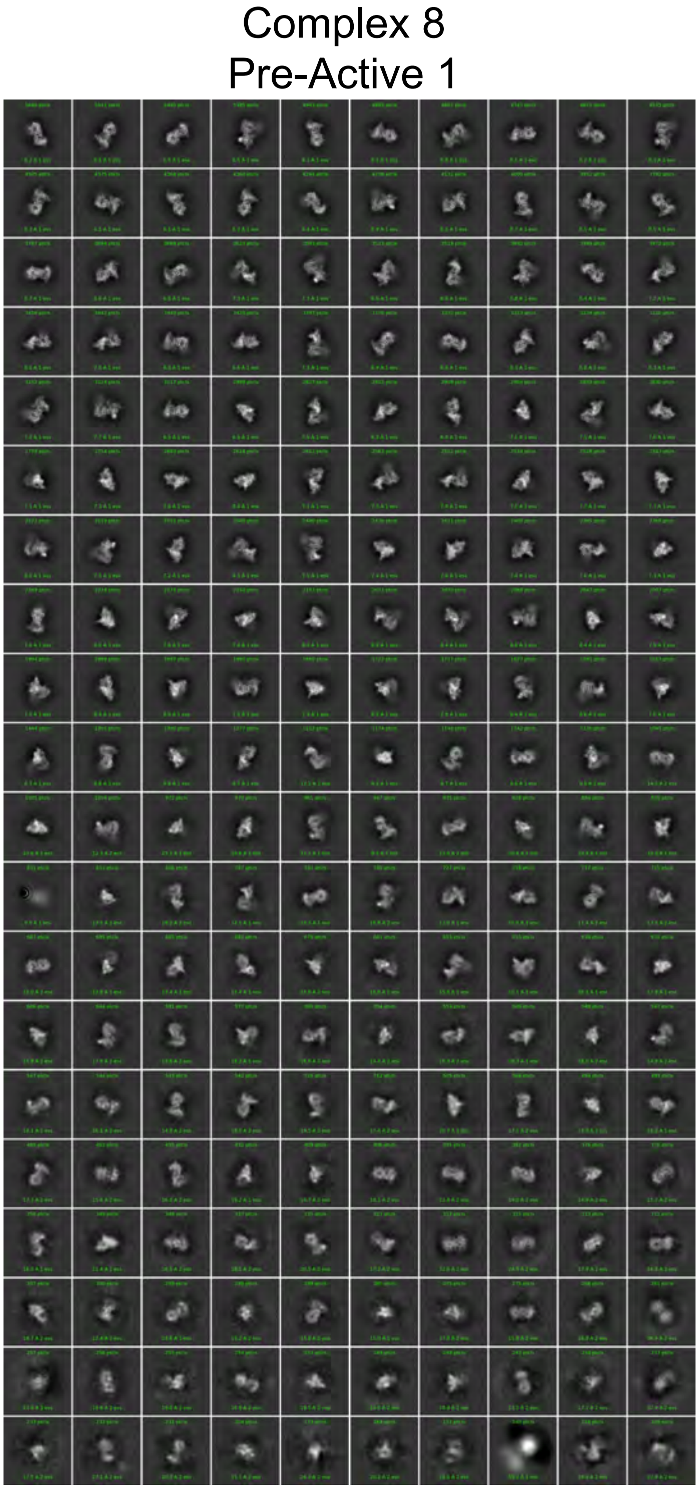

Figure S3, continued

Complex 9
Pre-Active 2

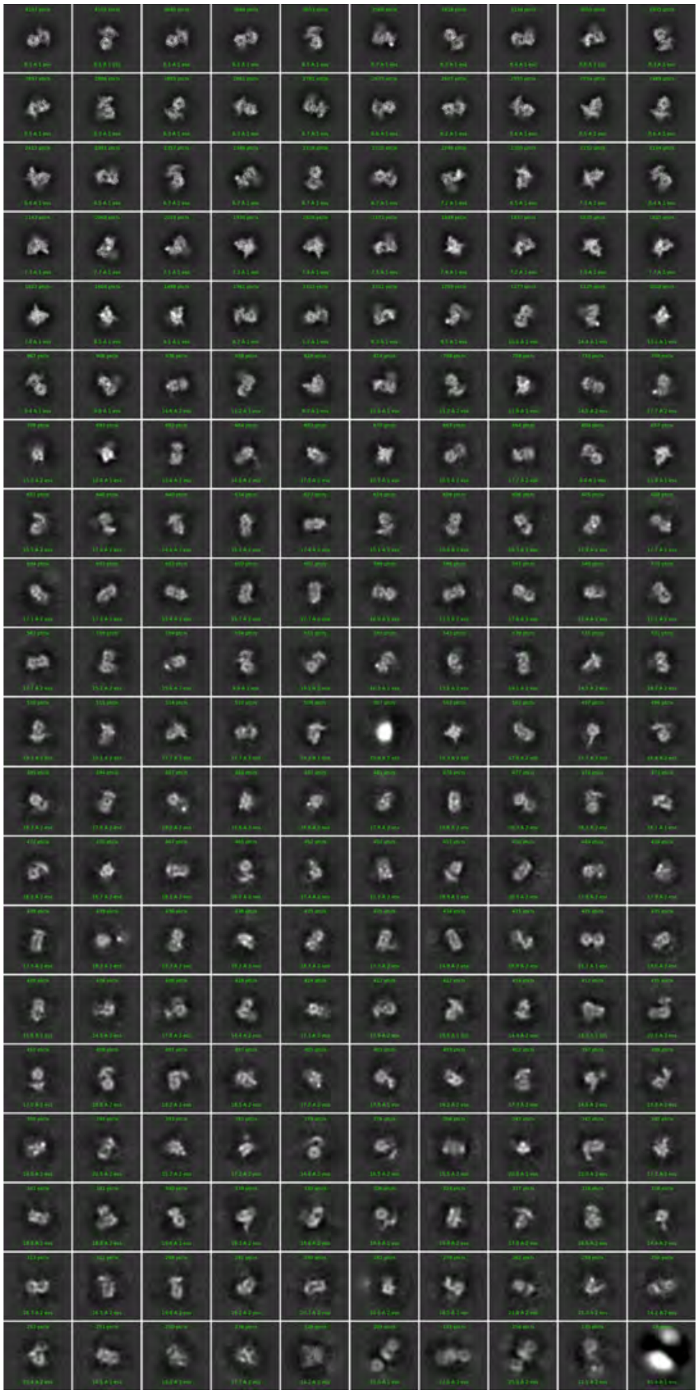

Figure S4

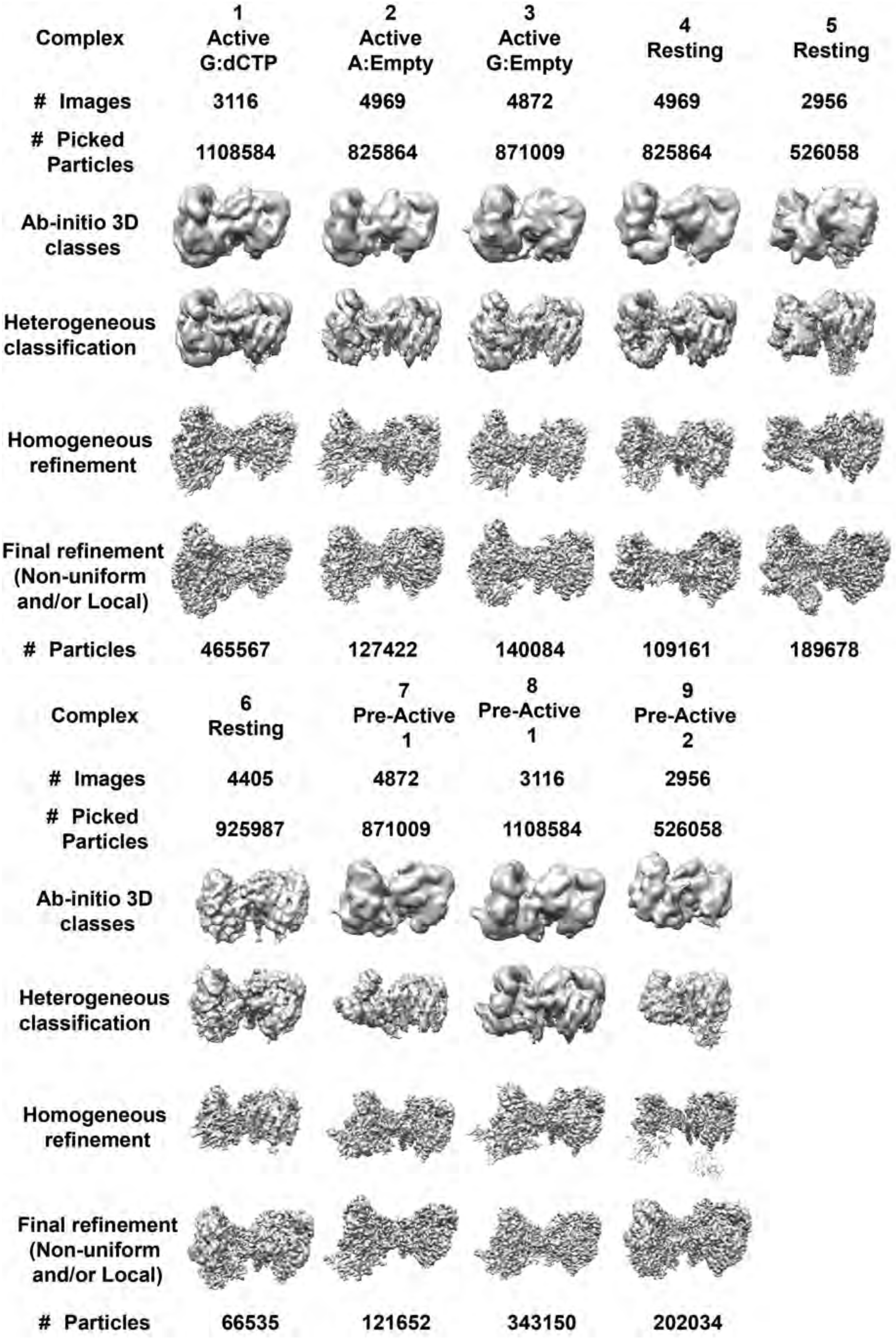


**Fig. S4. Cryo-EM Structure Determination.** Statistics and maps from stages of model building for nine bRT-Avd:RNA$^{\Delta 98}$ complexes (numbered as in Table S1).

Figure S5

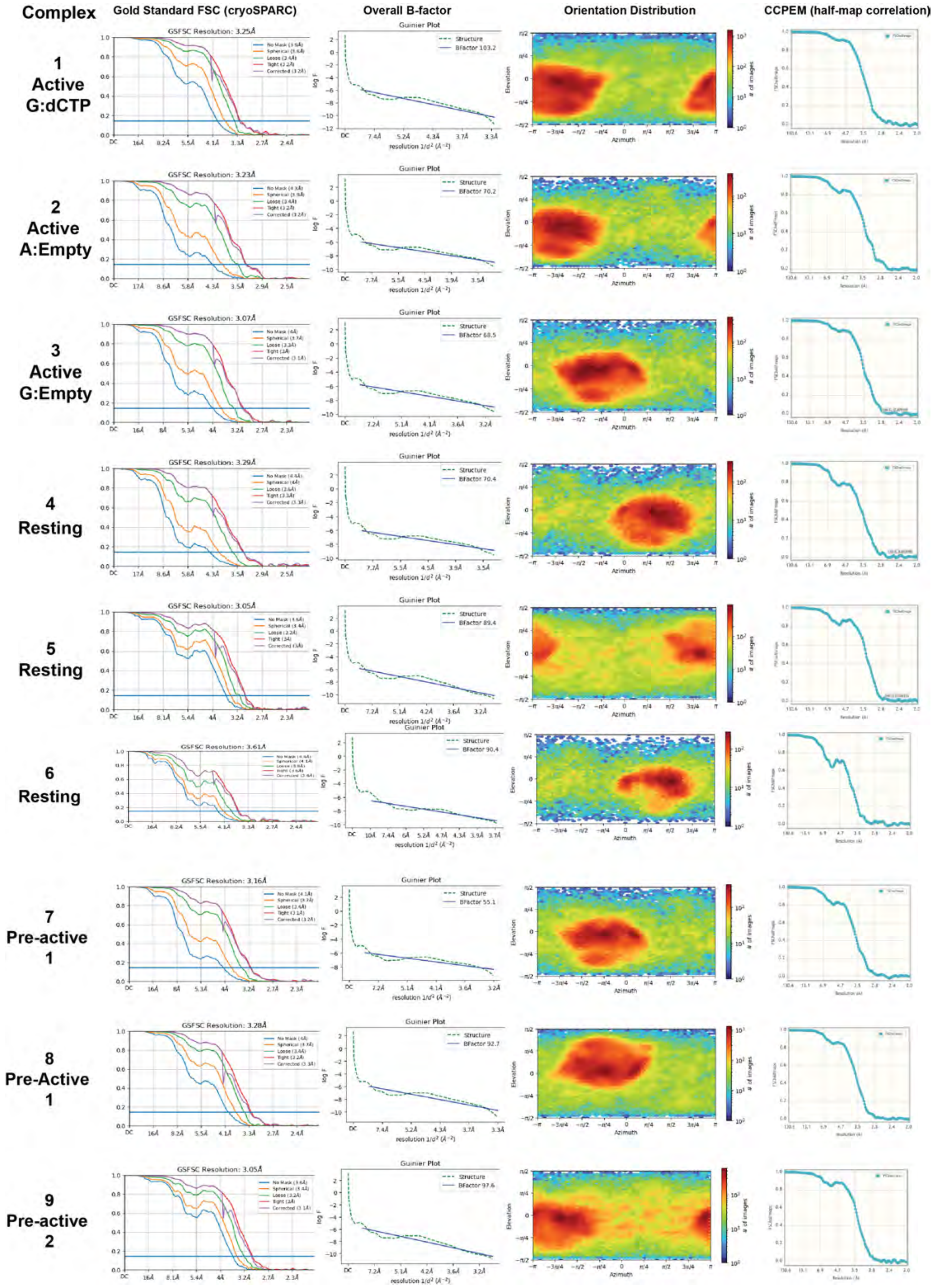
Complex
Gold Standard FSC (cryoSPARC)
Overall B-factor
Orientation Distribution
CCPEM (half-map correlation)
1 Active G:dCTP
GSFSC Resolution: 3.25Å
BFactor 103.2
2 Active A:Empty
GSFSC Resolution: 3.23Å
BFactor 70.2
3 Active G:Empty
GSFSC Resolution: 3.07Å
BFactor 68.5
4 Resting
GSFSC Resolution: 3.29Å
BFactor 70.4
5 Resting
GSFSC Resolution: 3.05Å
BFactor 89.4
6 Resting
GSFSC Resolution: 3.61Å
BFactor 90.4
7 Pre-active 1
GSFSC Resolution: 3.16Å
BFactor 55.1
8 Pre-Active 1
GSFSC Resolution: 3.28Å
BFactor 92.7
9 Pre-active 2
GSFSC Resolution: 3.05Å
BFactor 97.6
Guinier Plot
Structure
log F
resolution 1/d² (Å⁻²)
Elevation
Azimuth
### of images

**Fig. S5. Details of bRT-Avd:RNA$^{\Delta 98}$ complexes.** Sets of horizontal panels indicate cryoSPARC generated GSFSC curves, global B-factor of map, orientation distribution, and CCPEM derived half map correlation of nine bRT-Avd:RNA$^{\Delta 98}$ complexes (numbered as in Table S1).

Figure S6

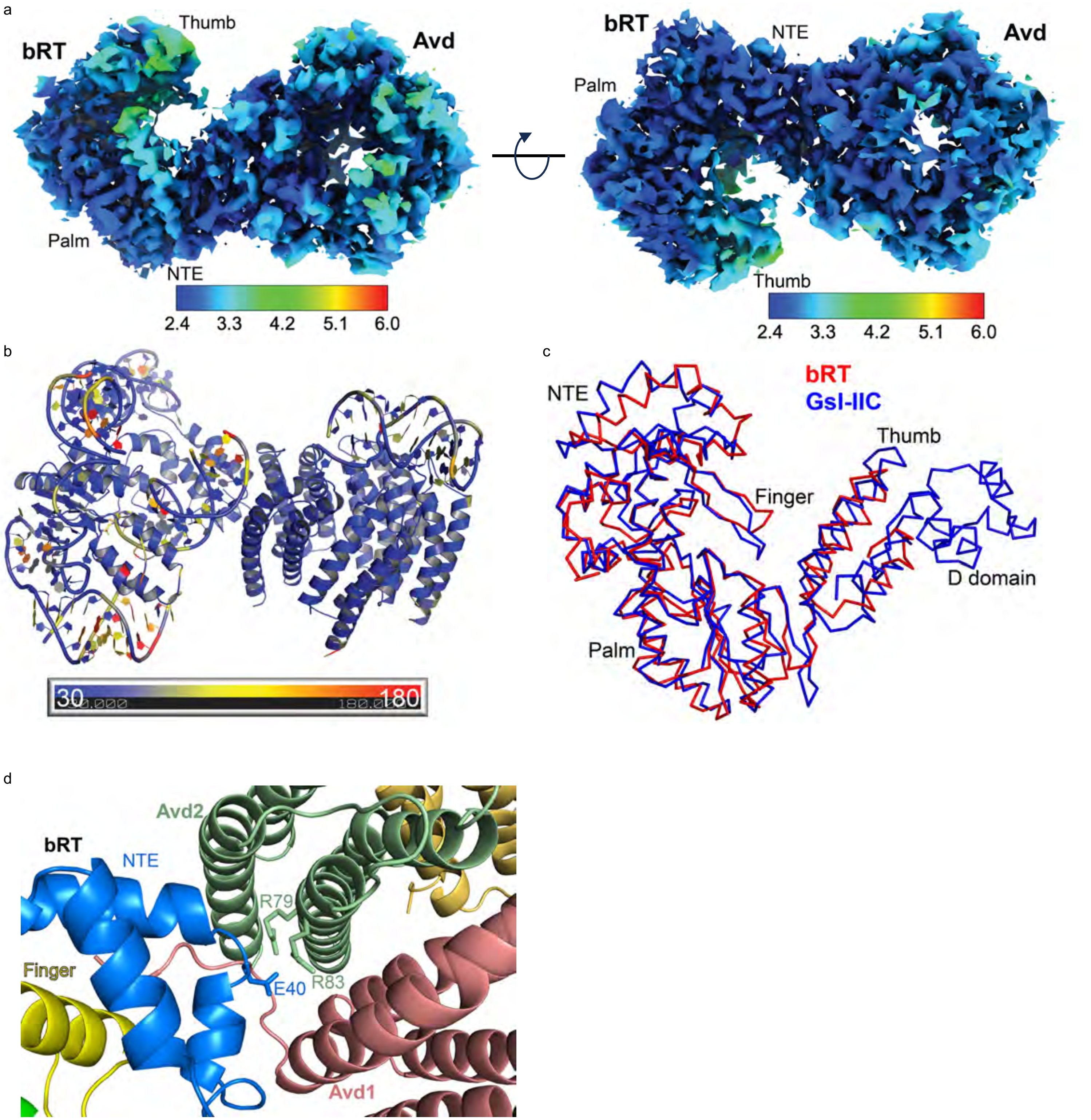


**Fig. S6. bRT-Avd. a.** Two views of local resolution map of bRT-Avd (left, in same orientation as Fig. 2a). Resolution scale in Å below. **b.** bRT-Avd:RNA$^{\Delta 98}$ colored by B-factors derived from Q-scores (blue-yellow-red, 30-105-180 $Å^2$). The resolution of bRT and Avd based on Q-scores was 2.6-2.9 Å, and that of the RNA 3.1 Å. **c.** bRT (red) superposed with the group II intron RT GsI-IIC (blue) in 'right-hand' view of polymerases. Subdomains of the RTs are indicated. GsI-IIC has a D domain that is not present in bRT or other DGR RTs. **d.** Interaction between bRT E40 and Avd R79 and R83. Avd R79A and R83A do not bind bRT and cause loss of function (*2*). Coloring as in Fig. 2a.

Figure S7

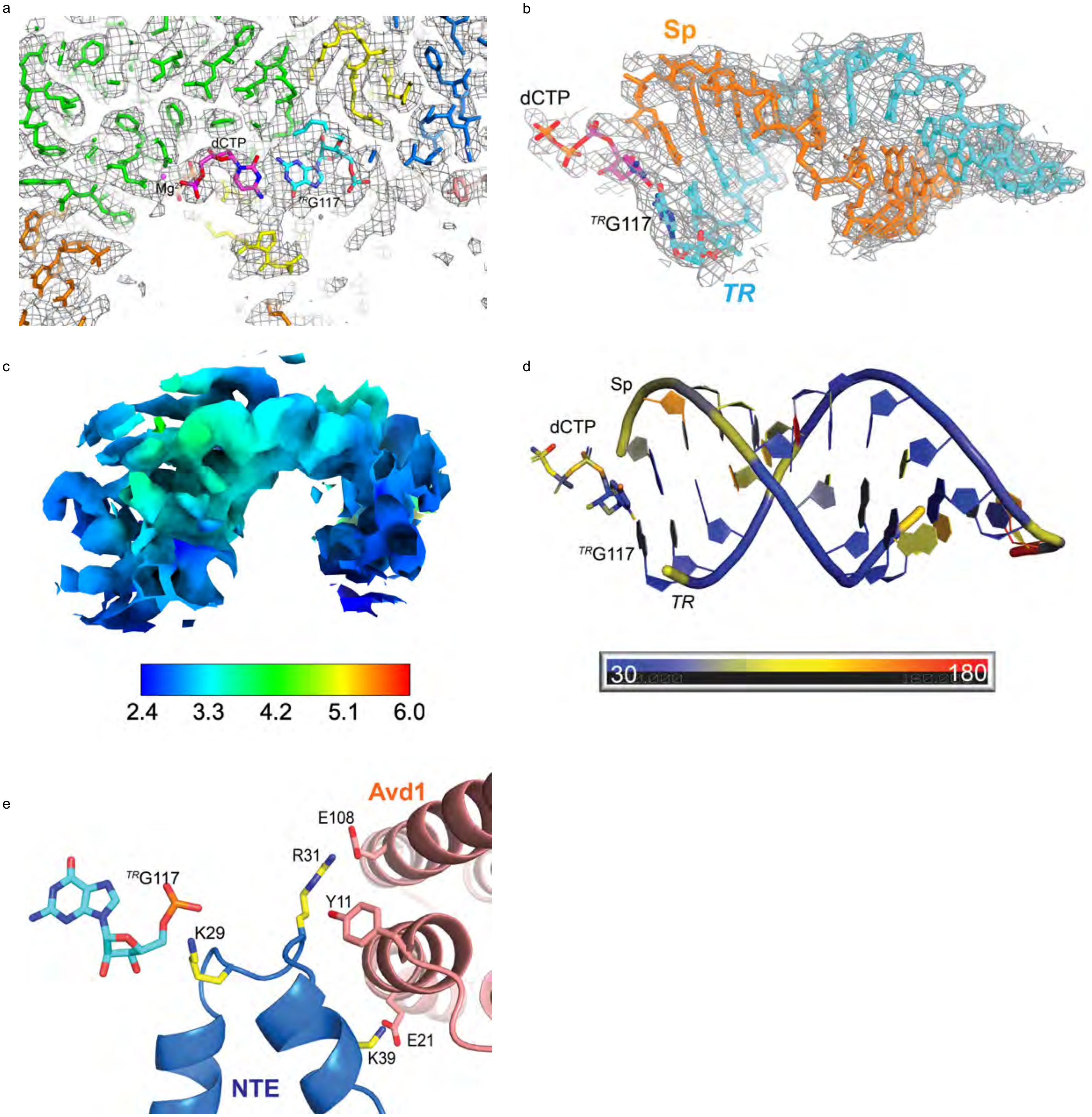


**Fig. S7. Template:Primer Duplex. a.** Cryo-EM density at the bRT active site, contoured at 6σ. Coloring is as in Fig. 2a in this and other panels. **b.** Cryo-EM density for the *TR*:Sp template:primer duplex, contoured at 6σ and within 2.5 Å of the duplex. **c.** Local resolution map of *TR:Sp* template:primer duplex. Resolution scale in Å below. **d.** *TR*:Sp duplex and dCTP colored by B-factors derived from Q-scores. The *TR*:Sp duplex (excluding unpaired $^{TR}$128 and 129) had a resolution of 3.1 Å based on the Q-score. **e.** Interaction of bRT NTE with $^{TR}$G117 and Avd. Interacting amino acids in bonds representation (carbon yellow for bRT, salmon for Avd, and cyan for $^{TR}$G117; oxygen red; nitrogen blue; and phosphorus orange).

Figure S8

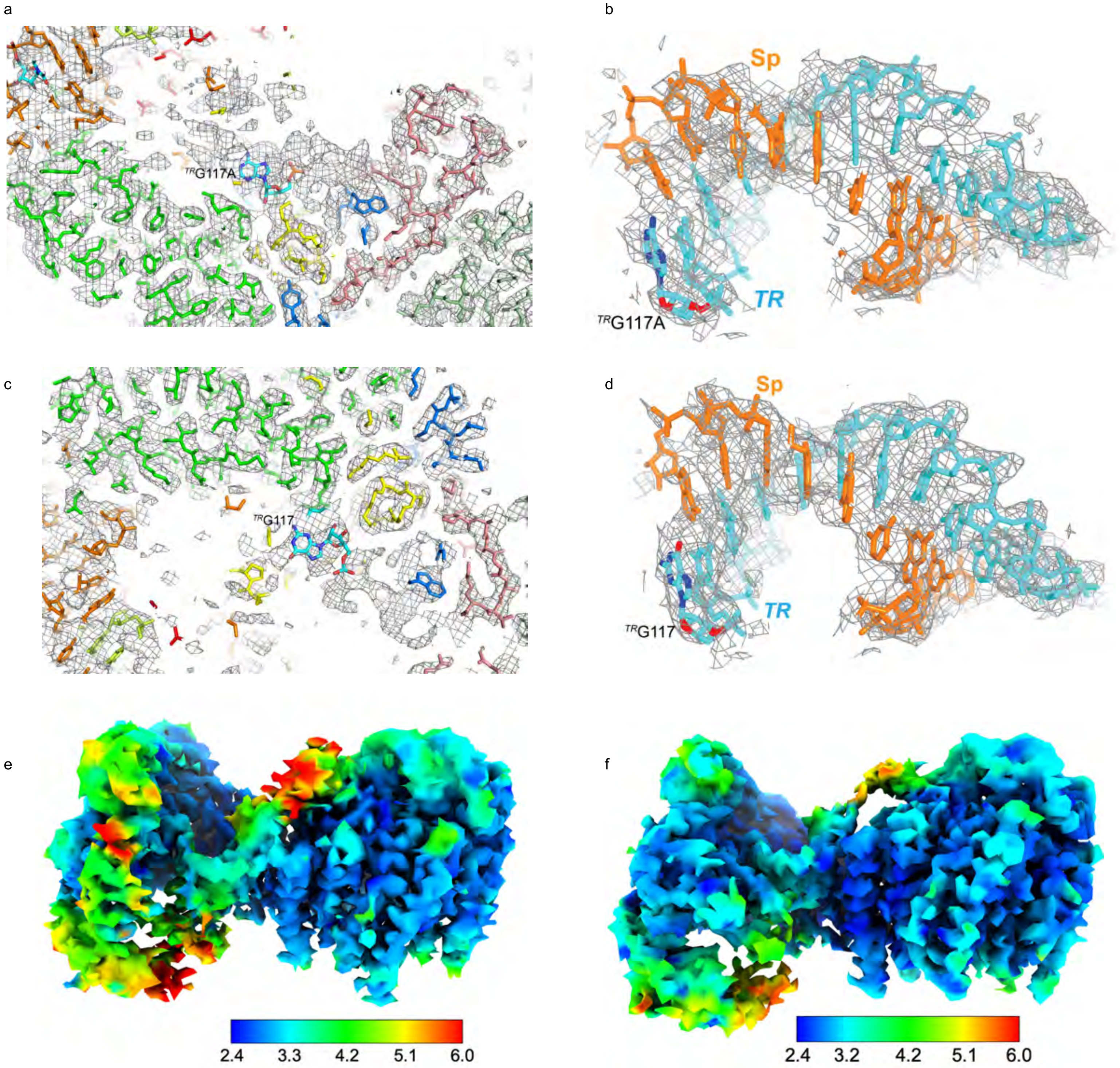


**Fig. S8. Active A:Empty and G:Empty. a and b.** Cryo-EM density at the bRT active site for the Active A:Empty RNP (a) and within 2.5 Å of the *TR:*Sp template:primer duplex (b), contoured at 6σ. Coloring is as in Fig. 2a in this and other panels. **c and d.** Cryo-EM density at the bRT active site for the Active G:Empty RNP (c) and within 2.5 Å of the *TR:*Sp template:primer duplex (d), contoured at 6σ. **e and f.** Local resolution maps of Active A:Empty (e, Table S1, Complex 2) and Active G:Empty RNPs (f, Table S1, Complex 3). Resolution scale in Å below.

Figure S8 continued

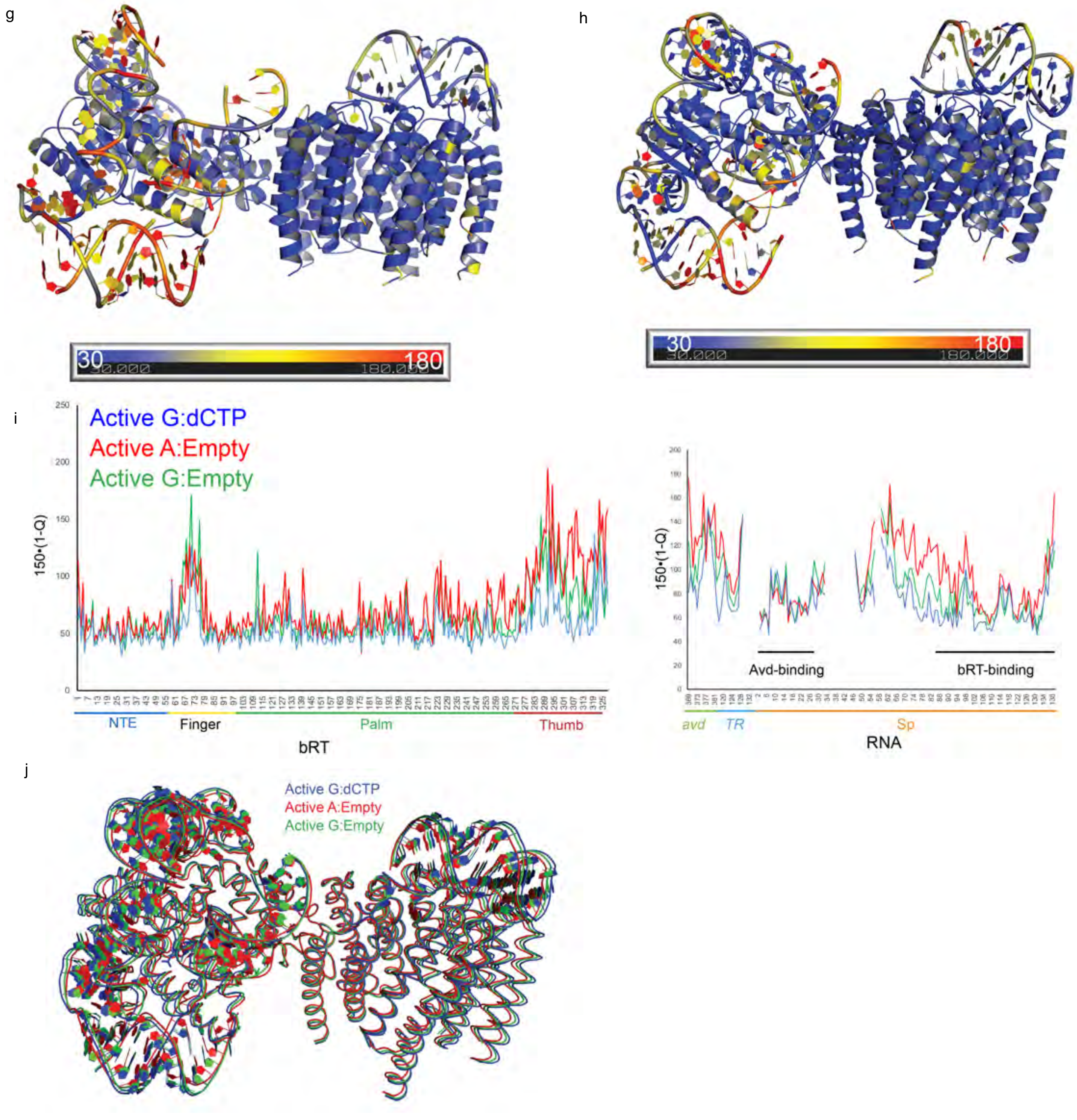


**Fig. S8 continued. g and h.** A:Empty RNP (g) and G:Empty (h) RNPs colored by B-factors derived from Q-scores. **i.** Plots of derived B-factor (B = 150 • (1-Q) ) as a function of bRT amino acid (left) or RNA nucleotide (right) in Active G:dCTP (blue, Complex 1), Active A:Empty (red, Complex 2), and Active G:Empty RNPs (green, Complex 3). **j.** Superposition of Active G:dCTP (blue, Complex 1), Active A:Empty (red, Complex 2), and Active G:Empty RNPs (green, Complex 3). RMSD of 1.6 Å for 10,218 atoms between Active G:dCTP and Active A:Empty RNPs, and 0.7 Å for 10,265 atoms between Active G:dCTP and Active A:Empty RNPs.

Figure S8 continued

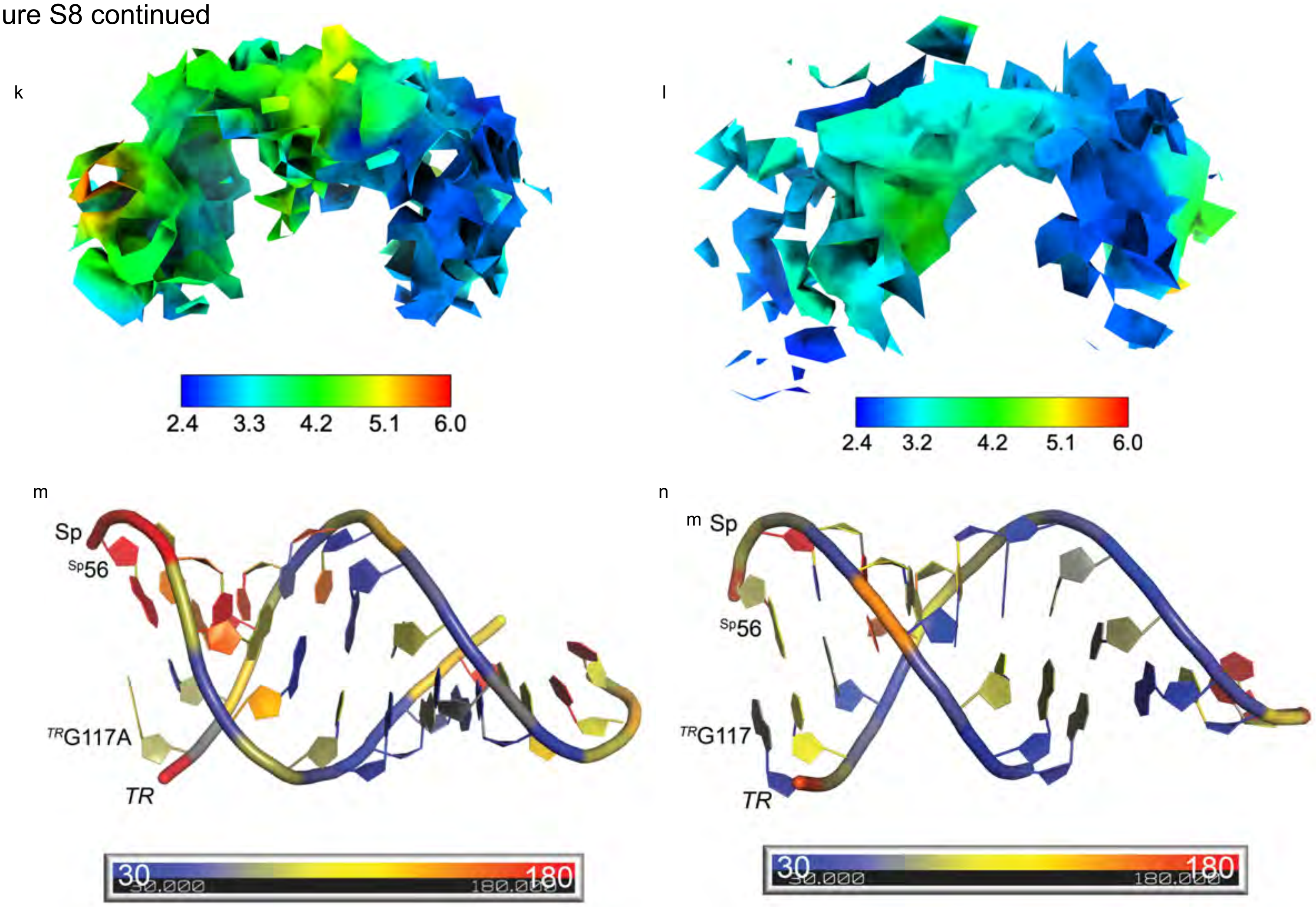


**Fig. S8 continued. k and l.** Local resolution of *TR*:Sp active site duplex in Active A:Empty (k) and Active G:Empty (l) RNPs. Resolution scale in Å below. **m and n.** *TR*:Sp duplex in the Active A:Empty (m) and Active G:Empty (n) RNPs colored by B-factors derived from Q-scores. The *TR*:Sp duplex (excluding unpaired $^{TR}$128 and 129) had a resolution of 3.8 and 4.2 Å based on the Q-score in A:Empty and G:Empty RNPs, respectively.

Figure S9

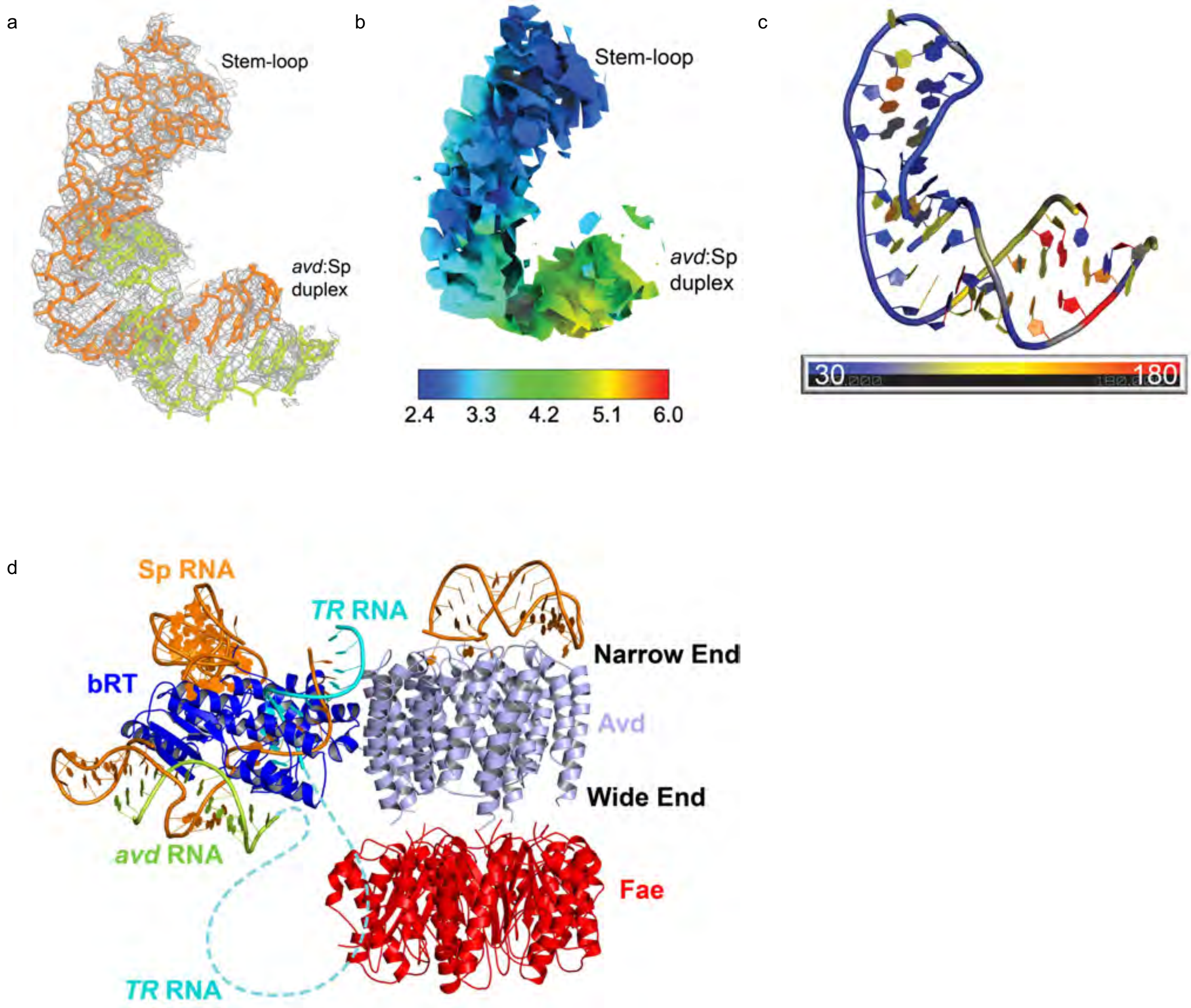


**Fig. S9. *c*PRT Stop. a.** Cryo-EM density within 2.5 Å of the *c*PRT Stop, contoured at 6σ. Coloring is as in Fig. 2a. **b.** Local resolution map of *c*PRT Stop. Resolution scale in Å below. **c.** *c*PRT Stop colored by B-factors derived from Q-scores. The *c*PRT duplex had a resolution of 4.1 Å based on the Q-score, and the stem-loop with unpaired nucleotides had a resolution of 2.9 Å. **d.** Model of bRT-Avd-Fae:RNA$^{\Delta 98}$ complex. Fae (red, PDB 1Y60) was placed at the wide end of the Avd barrel for visual purposes. No interpretable density exists for Fae. Coloring as in Fig. 1d. Dotted line indicates potential location of intact *TR*.

Figure S10

a

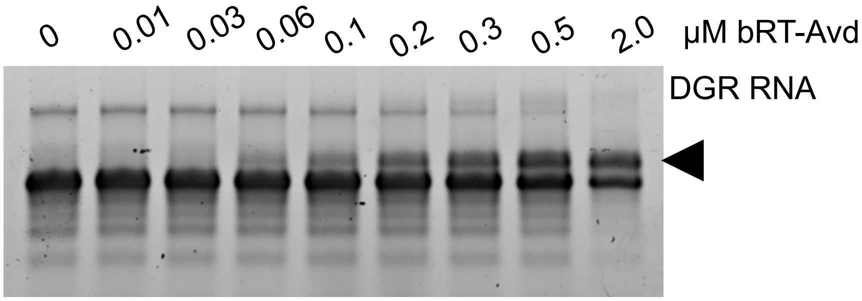


| Accession no. | -10lgP | Coverage (%) | Peptide no. | Unique peptide no. | Average Mass | Description |
|---|---|---|---|---|---|---|
| NP_958675.1 | 187.98 | 42 | 18 | 18 | 38035 | bRT |
| NP_958676.1 | 143.19 | 62 | 9 | 9 | 14472 | Avd |

b

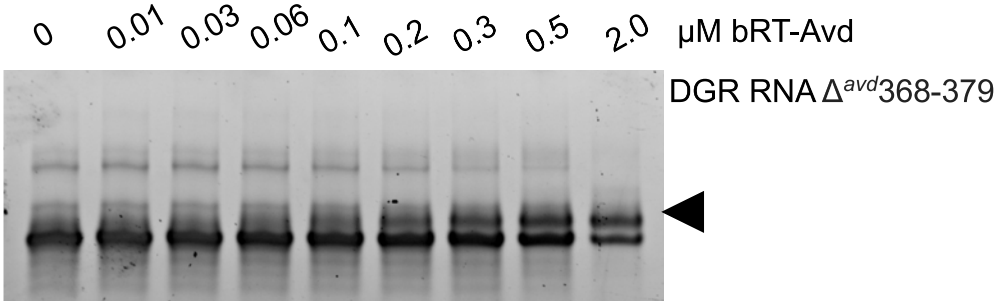


| Accession no. | -10lgP | Coverage (%) | Peptide no. | Unique peptide no. | Average Mass | Description |
|---|---|---|---|---|---|---|
| NP_958675.1 | 140.97 | 30 | 9 | 9 | 38035 | bRT |
| NP_958676.1 | 241.20 | 82 | 20 | 20 | 14472 | Avd |

c

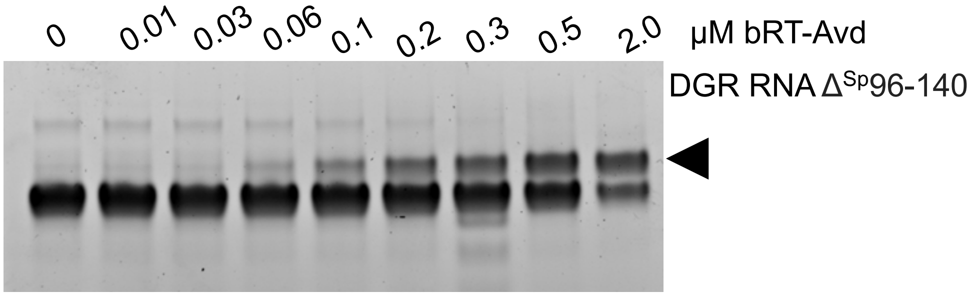


| Accession no. | -10lgP | Coverage (%) | Peptide no. | Unique peptide no. | Average Mass | Description |
|---|---|---|---|---|---|---|
| NP_958675.1 | - | - | - | - | 38035 | bRT |
| NP_958676.1 | 100.56 | 36 | 5 | 5 | 14472 | Avd |

d

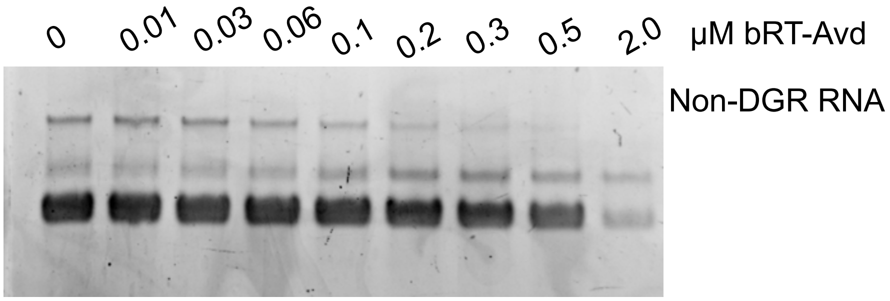


**Fig. S10. EMSA of wild-type and mutant DGR RNA.** Electrophoretic mobility shift of (**a**) DGR RNA, (**b**) DGR RNA Δ$^{avd}$368-379, (**c**) DGR RNA Δ$^{Sp}$96-140, and (**d**) non-DGR RNA by varying concentrations of bRT-Avd shown at top of each panel. Arrowheads indicate position of shifted bands. The bottom of each panel shows LC/MS-MS analysis of shifted EMSA bands. Lower quantities of RNA at higher bRT-Avd concentrations in the non-DGR RNA sample likely indicate non-specific aggregation of bRT-Avd with RNA. Representative gel from two independent replicates is shown.

Figure S11

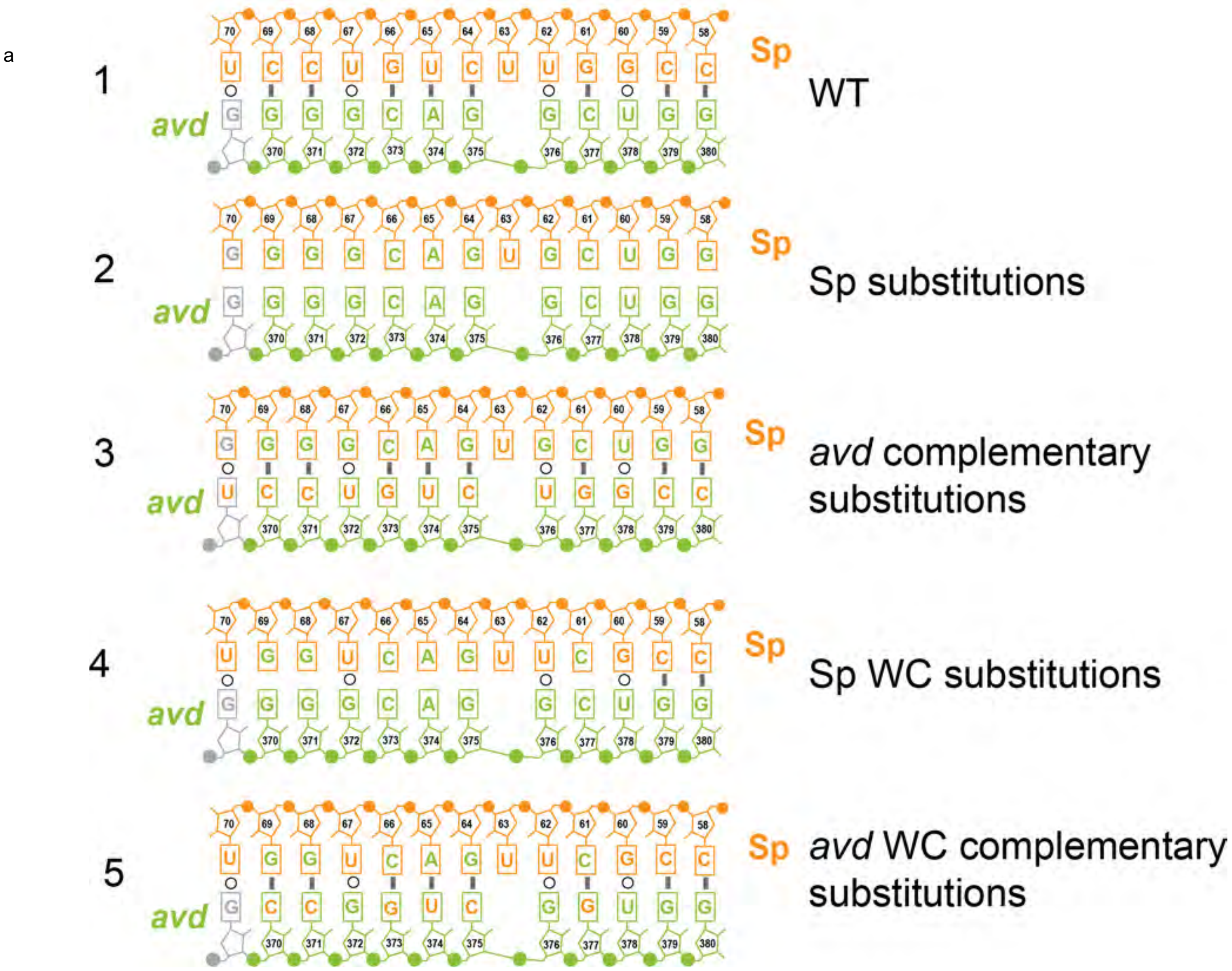


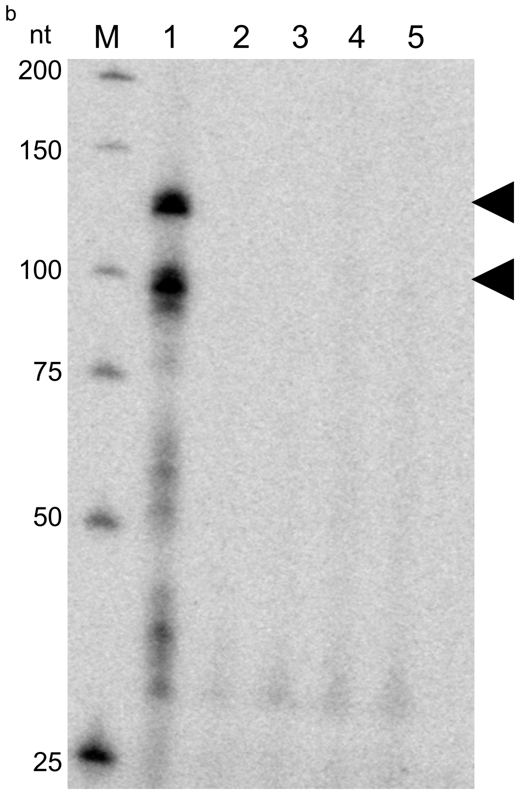


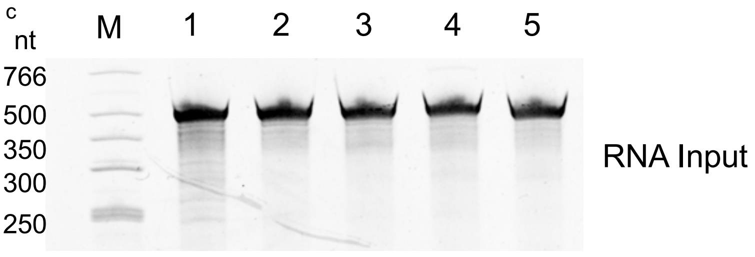


**Fig. S11. Disruption of *avd*-Sp base pairing in *c*PRT Stop. a.** Schematic of the 12 potential bps in the *avd*-Sp duplex (1, WT); (2) substitutions in Sp that disrupt the 12 bps; (3) complementary substitutions in *avd* that restore the 12 bps; (4) substitutions in Sp that disrupt six Watson-Crick bps; and (5) complementary substitutions in *avd* that restore the six Watson-Crick bps. Solid bars are Watson-Crick base pairs, and open circles wobble base pairs. **b**. cDNA synthesis by bRT-Avd with (1) wild-type or (2-5) mutant DGR RNA. The lane numbers correspond to the numbering of sequences in panel a. Arrowheads indicate the positions of ~90- and ~120-nt cDNAs. A representative gel from three independent replicates is shown. **c.** Input RNA for cDNA synthesis. The numbers correspond to the numbering of sequences depicted in panel a.

Figure S12

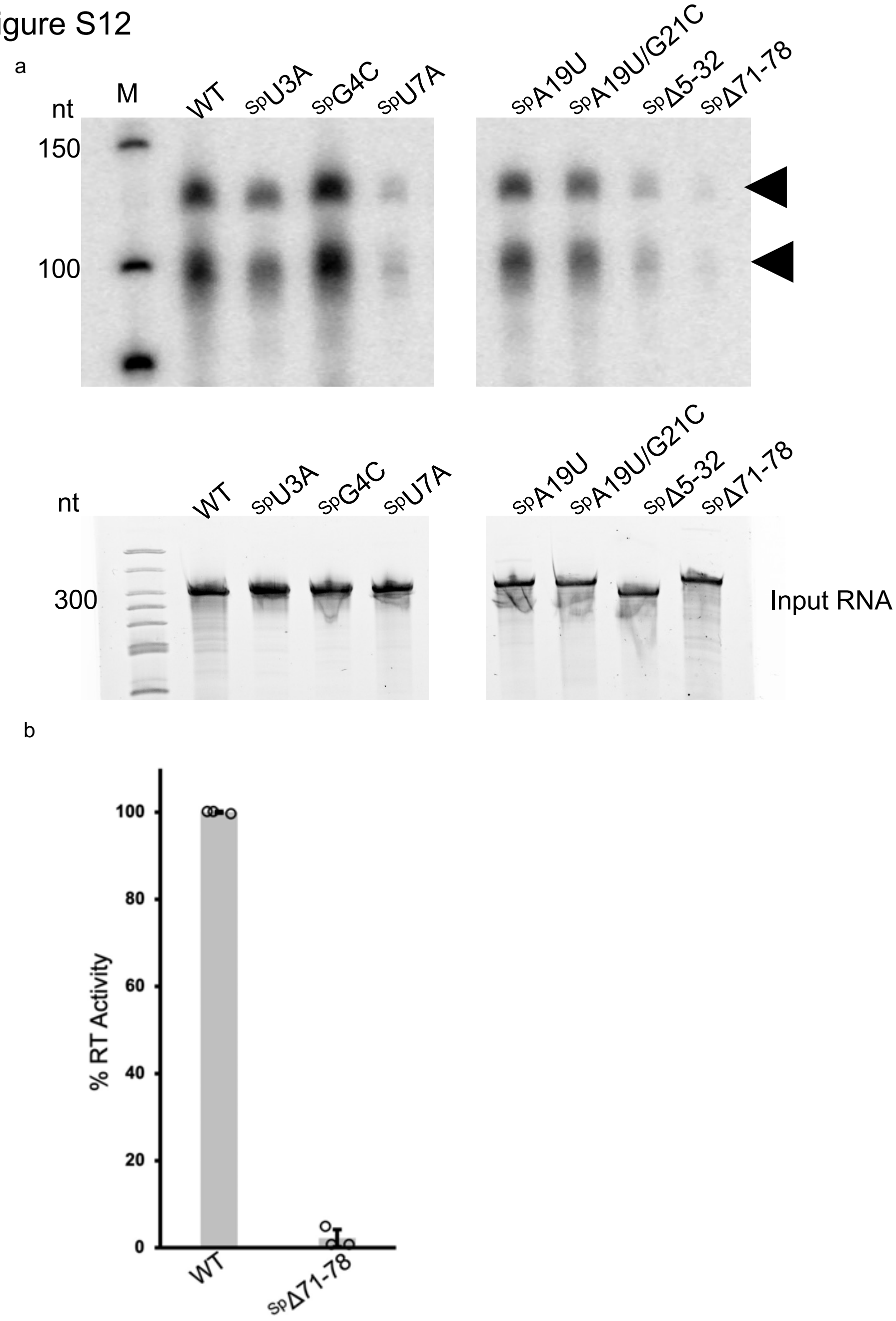


**Fig. S12. *c*PRT with DGR RNAs mutated in *c*PRT Stop or Avd-binding element. a.** cDNAs synthesized by bRT-Avd from wild-type or mutated DGR RNAs shown at top of each panel. Arrowheads correspond to cDNAs. Bottom of each panel shows input RNAs for cDNA synthesis. A representative gel from three independent replicates shown. An irrelevant lane is blanked out from the gels. **b.** *c*PRT activity of mutant DGR RNA relative to wild-type DGR RNA. Data are from experiments that were repeated three independent times, and means and standard deviations shown.

### Figure S13

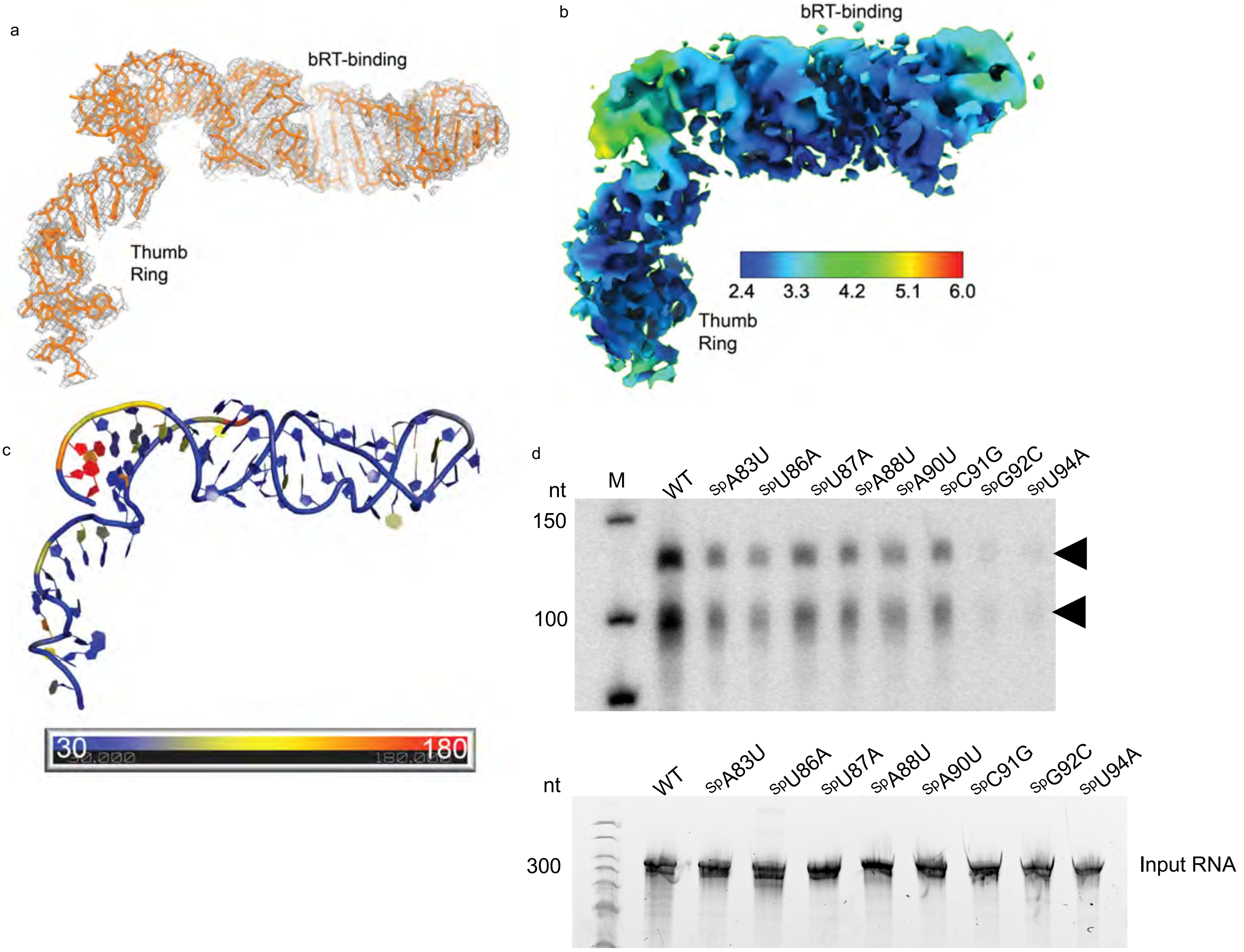


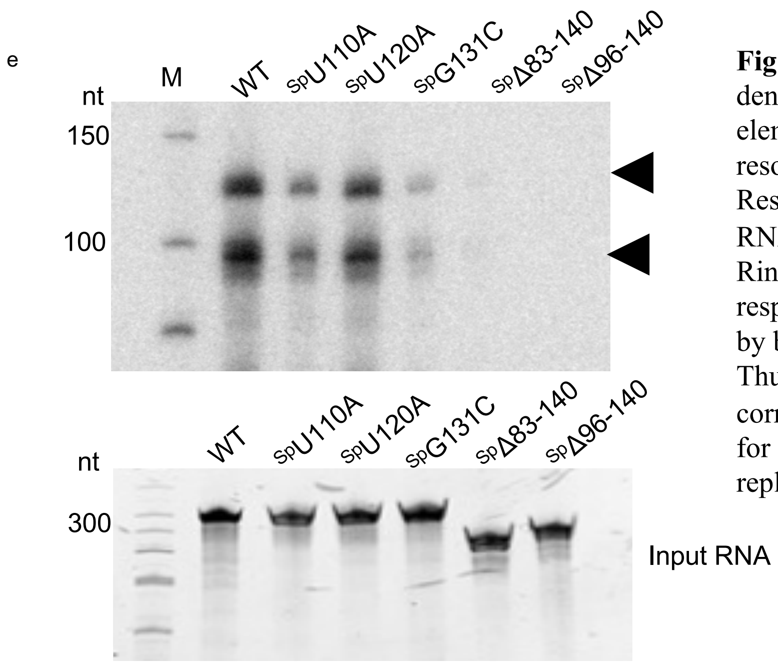


**Fig. S13. Thumb Ring and bRT-binding RNA. a.** Cryo-EM density within 2.5 Å of the Thumb Ring and bRT-binding RNA element, contoured at 6σ. Coloring is as in Fig. 2a. **b.** Local resolution map of Thumb Ring and bRT-binding RNA. Resolution scale in Å below. **c.** Thumb Ring and bRT-binding RNA colored by B-factors derived from Q-scores. The Thumb Ring and bRT-binding RNA had resolutions of 2.3 and 2.8 Å, respectively, based on the Q-score. **d and e.** cDNAs synthesized by bRT-Avd from wild-type or DGR RNAs mutated in the Thumb Ring (d) or bRT-binding RNA (e). Arrowheads correspond to cDNAs. Bottom of each panel shows input RNAs for cDNA synthesis. A representative gel from three independent replicates shown.

Figure S14

a

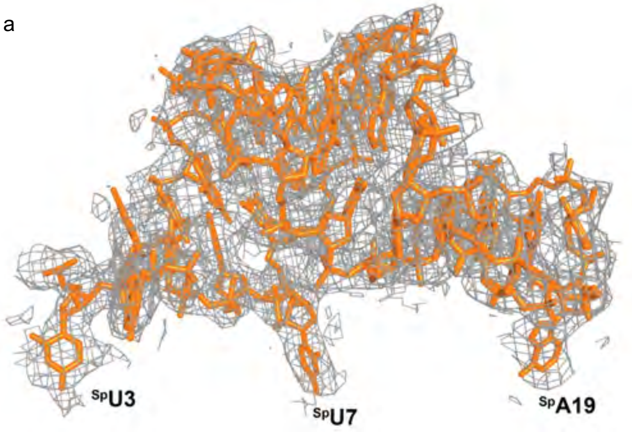


b

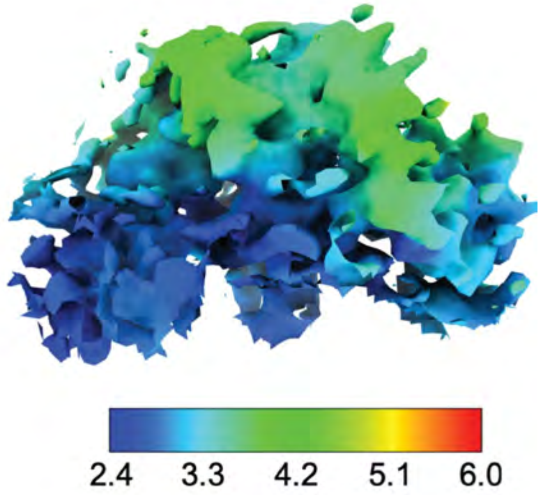


c

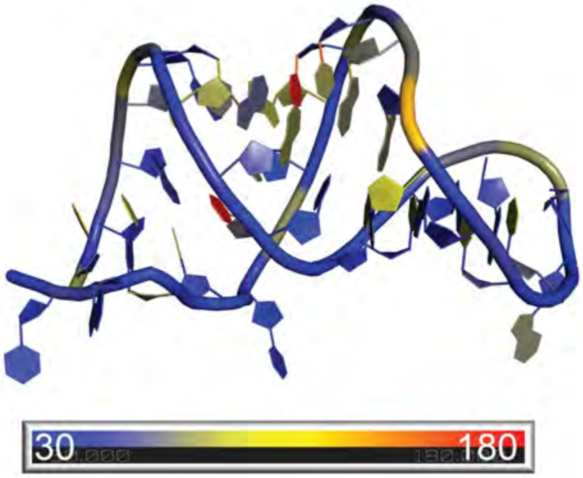


d

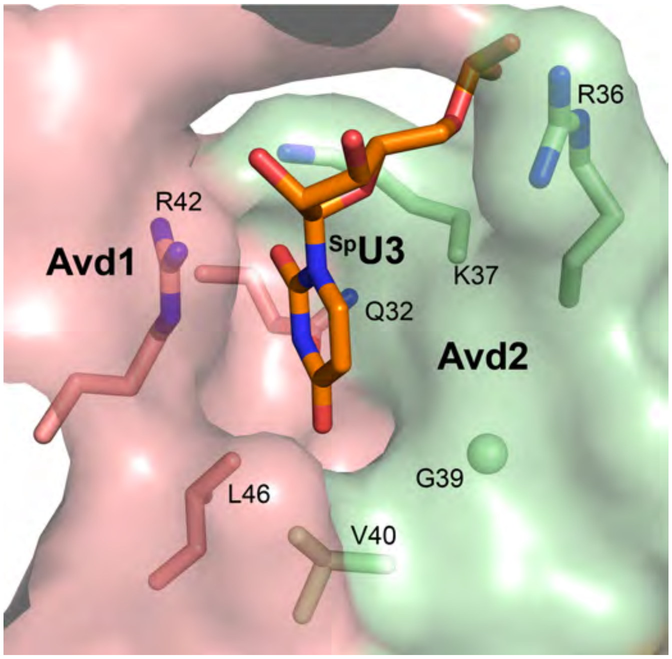


e

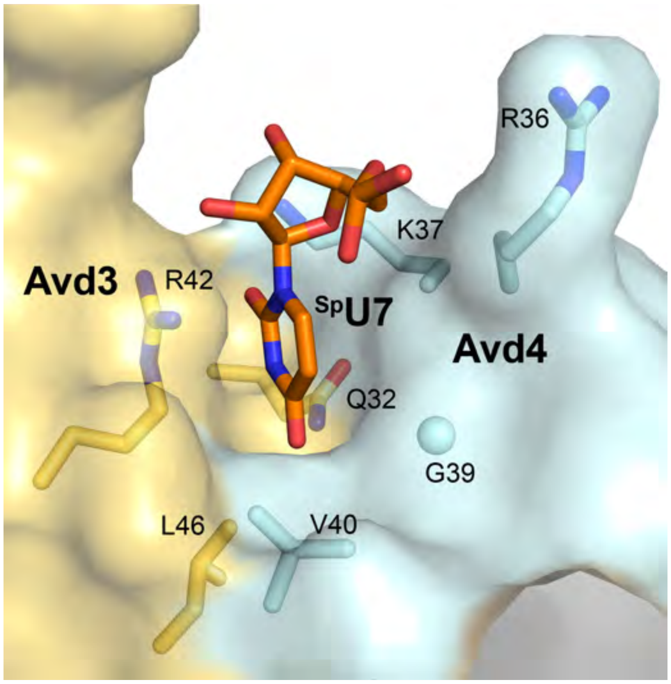


**Fig. S14. Avd-binding RNA. a.** Cryo-EM density within 2.5 Å of the Avd-binding RNA, contoured at 6σ. Coloring is as in Fig. 2a. **b.** Local resolution map of Avd-binding RNA. Resolution scale in Å below. **c.** Avd-binding RNA colored by B-factors derived from Q-scores. The Avd-binding RNA had a resolution of 3.0 Å based on the Q-score. **d.** Binding of flipped-out base $^{Sp}$U3 to a crevice between Avd1 and Avd2. **e.** Binding of flipped-out base $^{Sp}$U7 to a crevice between Avd3 and Avd4.

Figure S15

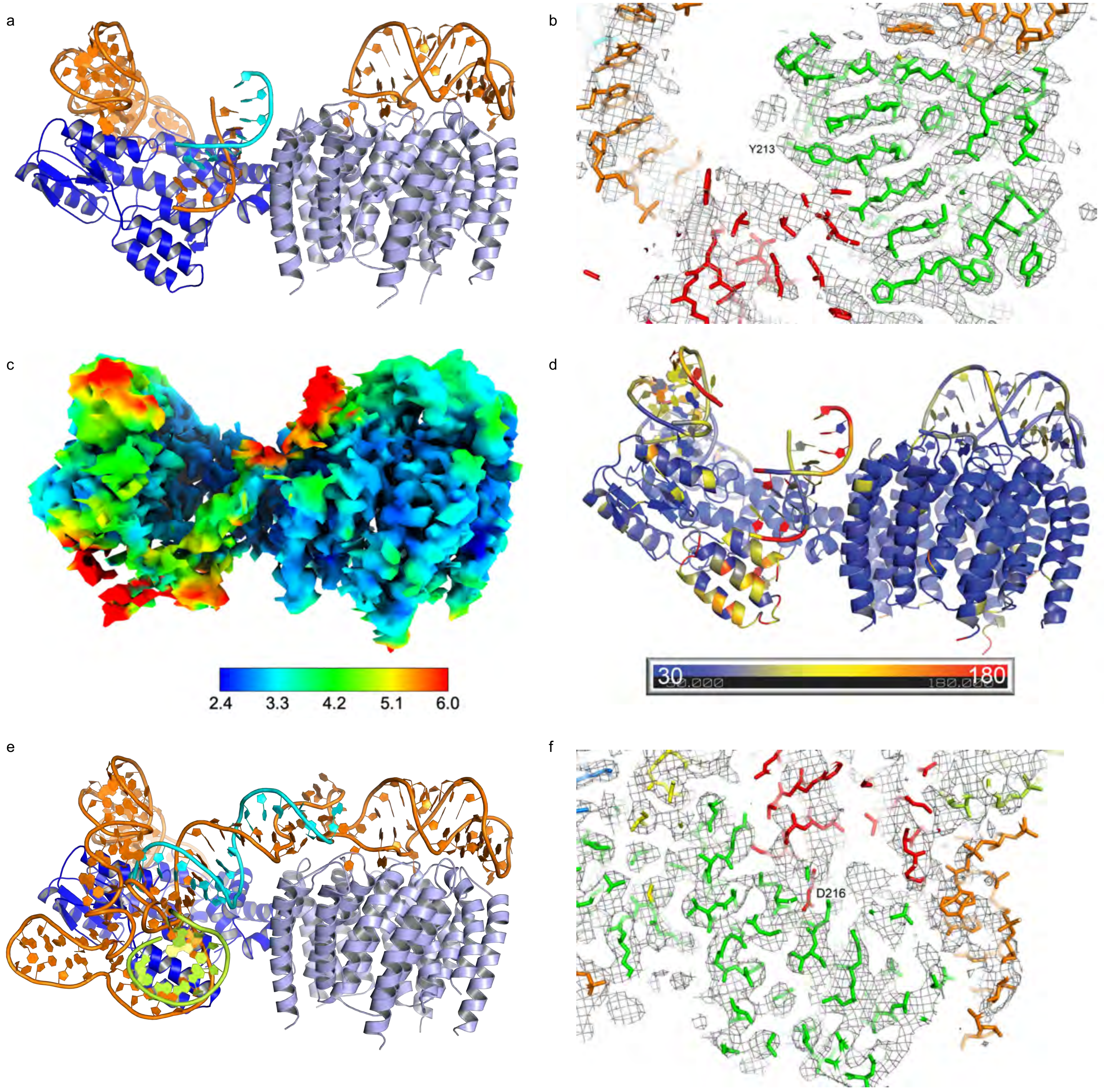


**Fig. S15. Resting Conformation. a.** bRT-Avd:RNA$^{\Delta 98}$ in Resting conformation (Complex 4; coloring as in Fig. 1d). **b.** Cryo-EM density for Complex 4 contoured at 6σ. Coloring is as in Fig. 2a. **c.** Local resolution map of Complex 4. Resolution scale in Å below. **d.** Complex 4 colored by B-factors derived from Q-scores. **e.** Complex 5, coloring as in Fig. 1d. **f.** Cryo-EM density for Complex 5 contoured at 6σ. Coloring as in Fig. 2a.

Figure S15 Continued

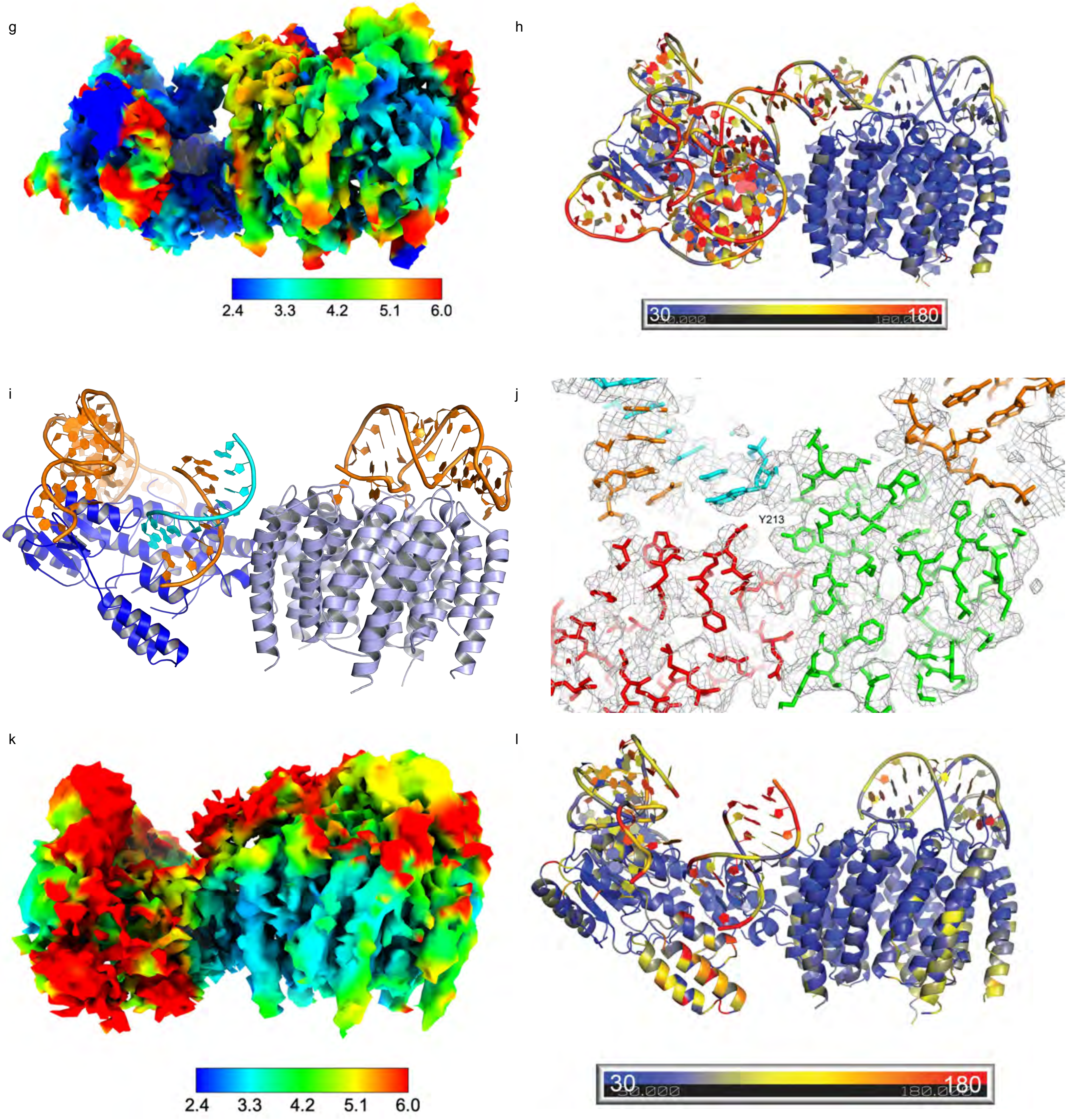


**Fig. S15 continued. g.** Local resolution map of Complex 5. Resolution scale in Å below. **h.** Complex 5 colored by B-factors derived from Q-scores. **i.** Complex 6, coloring as in Fig. 1d. **j.** Cryo-EM density for Complex 6, contoured at 6σ. Coloring is as in Fig. 2a. **k.** Local resolution map of Complex 6. Resolution scale in Å below. **l.** Complex 6 colored by B-factors derived from Q-scores.

Figure S16

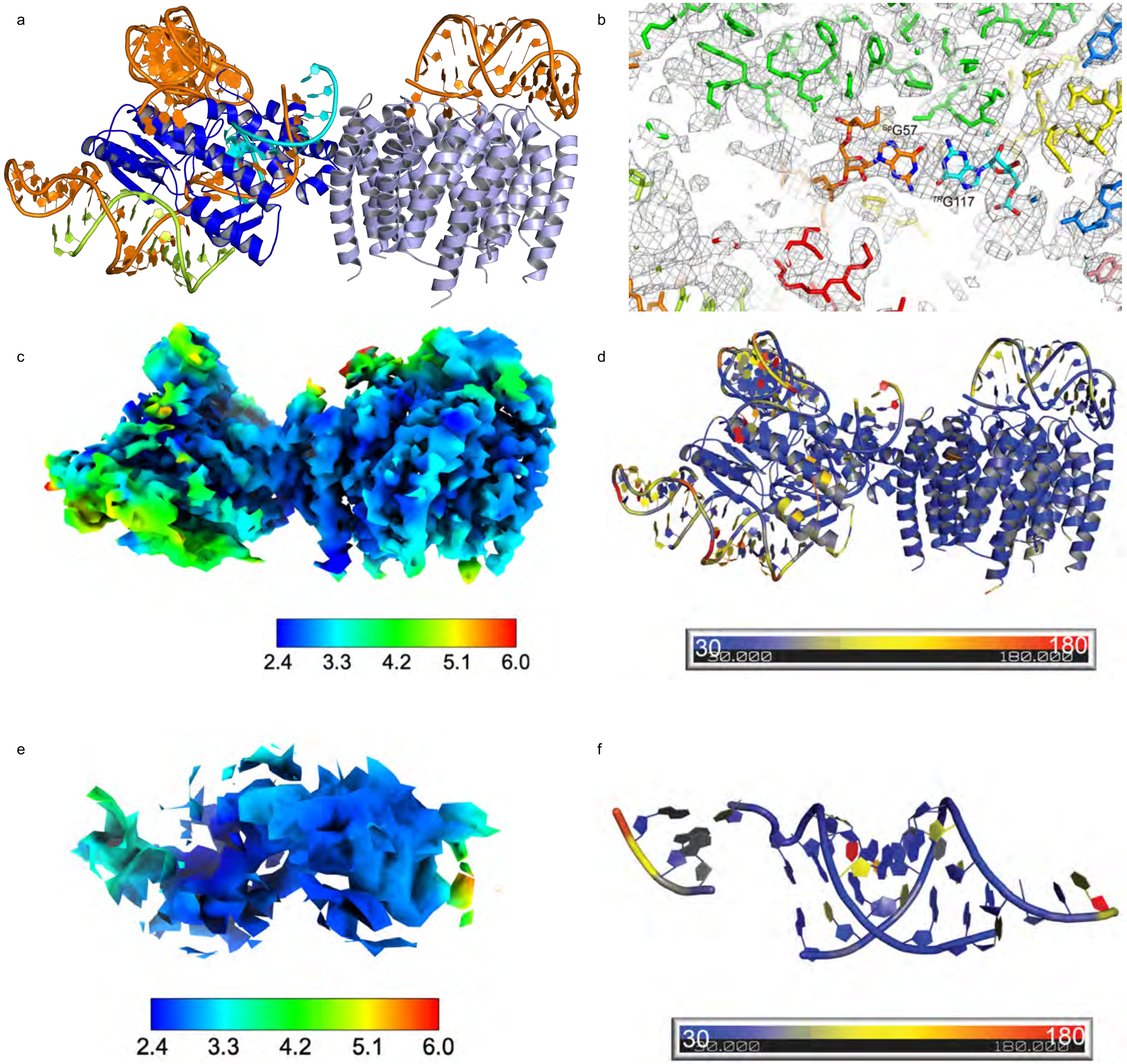


**Fig. S16. Pre-Active Conformation 1. a.** bRT-Avd:$RNA^{\Delta98}$ in Pre-Active 1 conformation (Complex 7; coloring as in Fig. 1d). **b.** Cryo-EM density for Complex 7 contoured at 6σ. Coloring is as in Fig. 2a. **c.** Local resolution map of Complex 7. Resolution scale in Å below. **d.** Complex 7 colored by B-factors derived from Q-scores. **e.** Local resolution map of *TR*:Sp duplex, including $^{Sp}$57 and 58 and $^{avd}$380-382, in Complex 7. Resolution scale in Å below. **f.** *TR*:Sp duplex, including $^{Sp}$57 and 58 and $^{avd}$380-382, in Complex 7 colored by B-factors derived from Q-scores.

Figure S16 continued

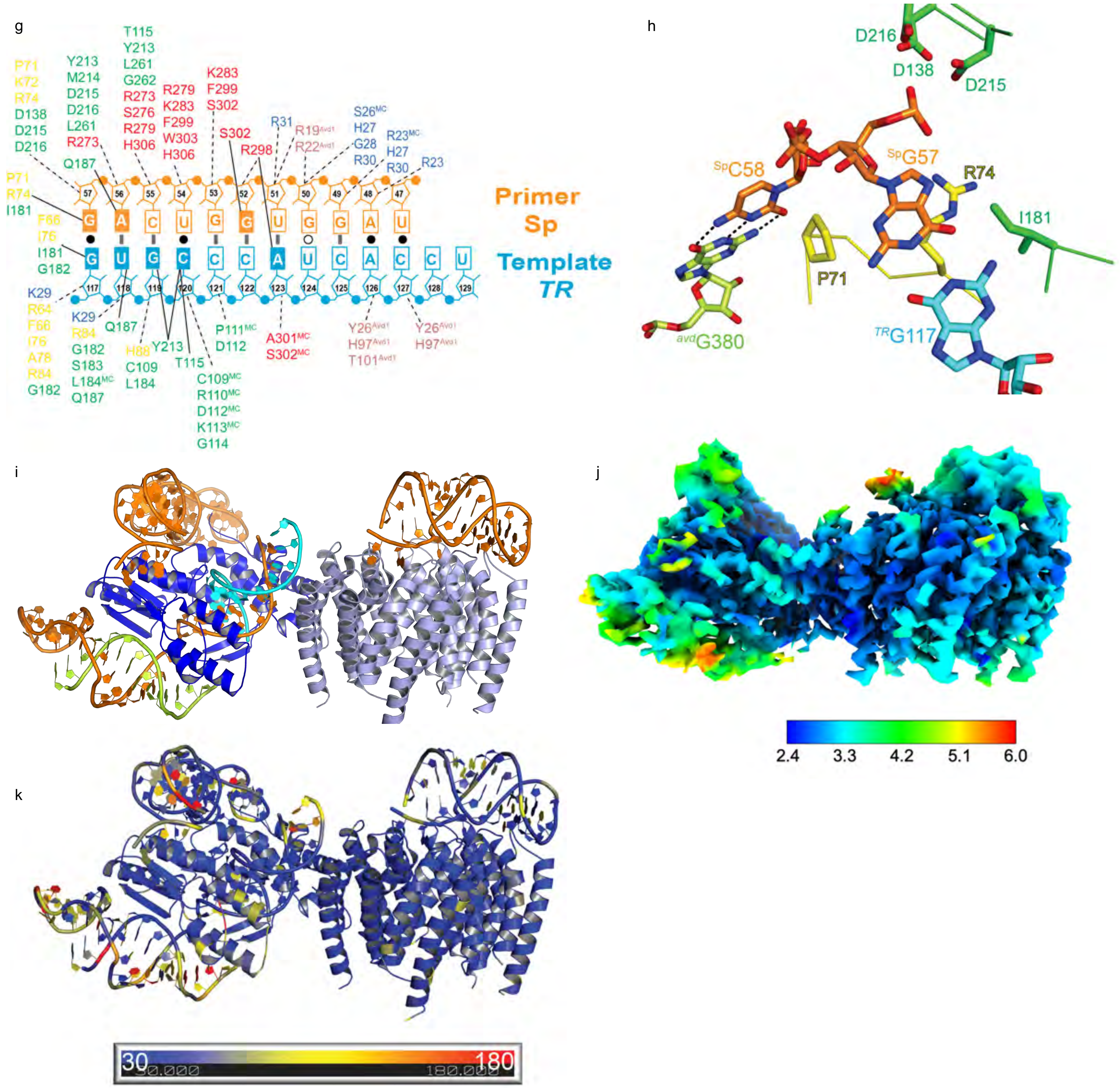


**Figure S16 continued. g.** Schematic of *TR*:Sp duplex in Complex 7, depicted as in Fig. 2c. **h.** Details of interactions of $^{Sp}$G57 and C58 with $^{TR}$G117 and $^{avd}$G380, respectively. bRT amino acids that contact $^{Sp}$G57 are shown. Coloring as in Fig. 2b, with oxygen red and nitrogen blue in the RNA. **i.** Complex 8, coloring as in Fig. 2a. **j.** Local resolution map of Complex 8. Resolution scale in Å below. **k.** Complex 8 colored by B-factors derived from Q-scores.

Figure S17

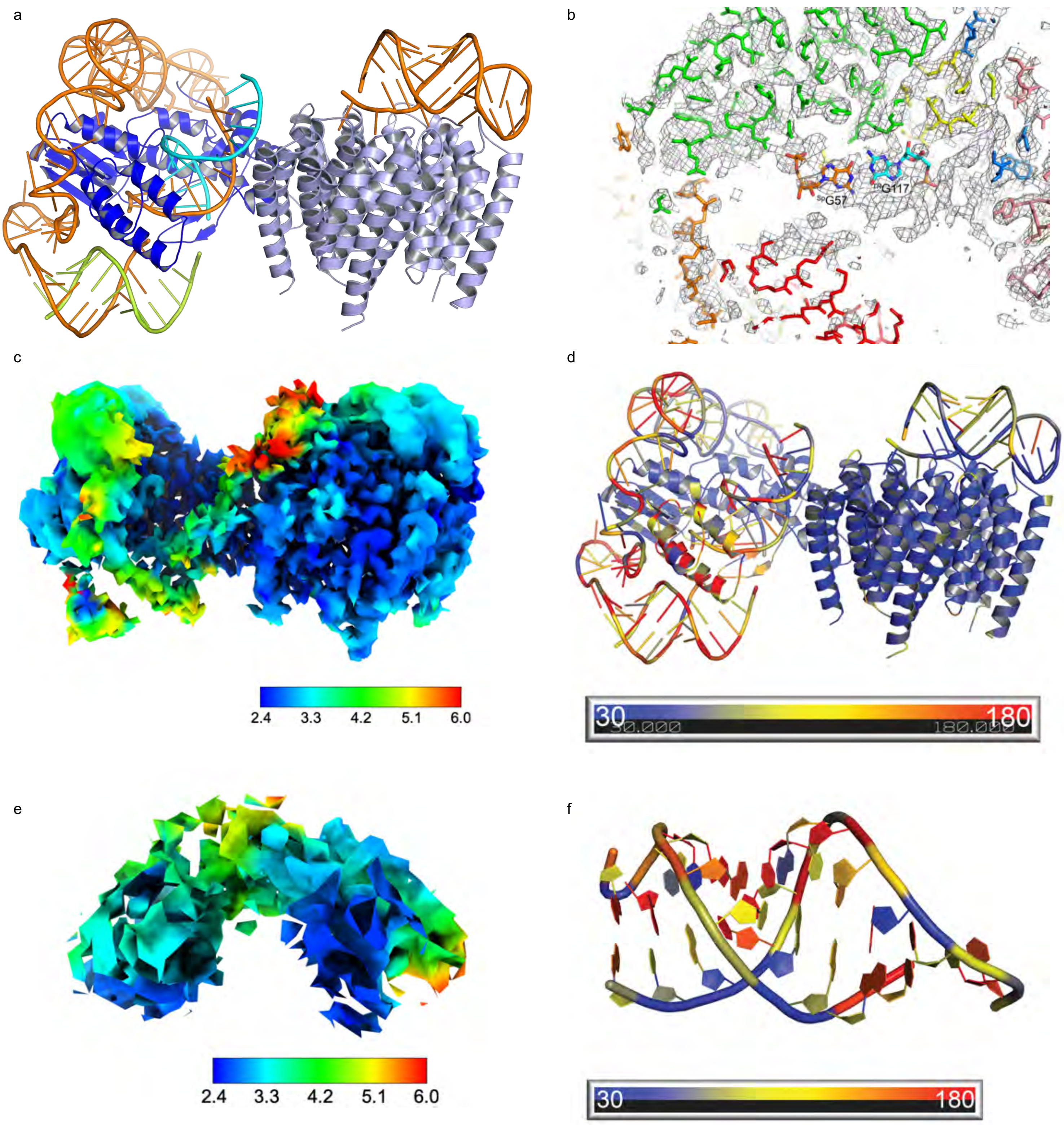


**Fig. S17. Pre-Active Conformation 2. a.** bRT-Avd:$RNA^{\Delta98}$ in Pre-Active 2 conformation (Complex 9; coloring as in Fig. 1d). **b.** Cryo-EM density for Complex 9 contoured at $6\sigma$. Coloring is as in Fig. 2a. **c.** Local resolution map of Complex 9. Resolution scale in Å below. **d.** Complex 9 colored by B-factors derived from Q-scores. **e.** Local resolution map of *TR*:Sp duplex in Complex 9. Resolution scale in Å below. **f.** *TR*:Sp duplex in Complex 9 colored by B-factors derived from Q-scores.

Figure S18

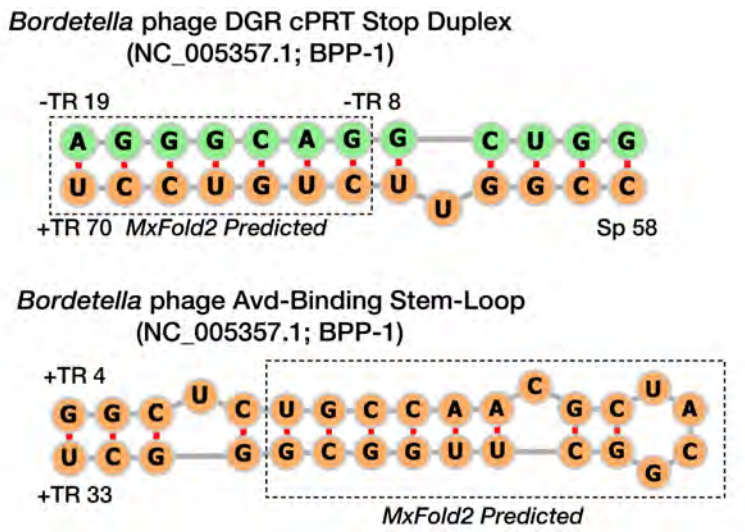


**Fig. S18. RNA structure prediction.** RNA structural features of the BΦ DGR. The *c*PRT Stop duplex and Avd-binding stem-loop elements determined by cryo-EM were compared with structural predictions from MxFold2 (*3*). The sequence upstream (i.e., *avd*) and downstream (i.e., Sp) of *TR* are in green and orange, respectively.

Figure S19

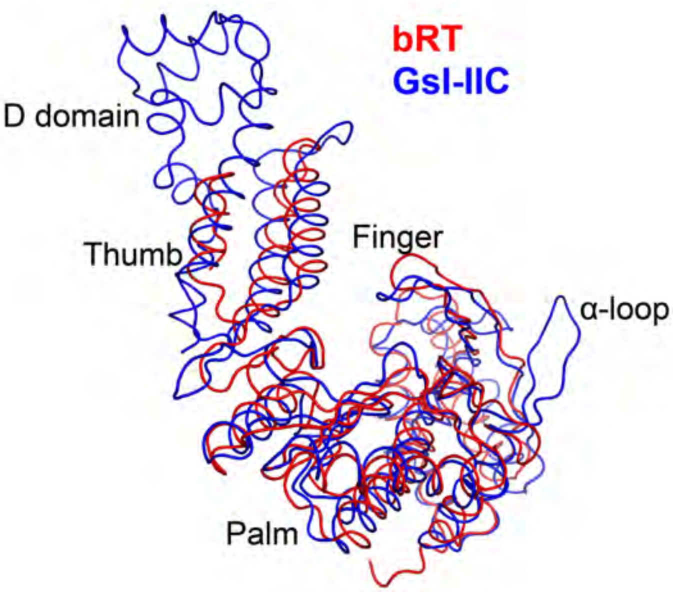


**Fig. S19. α-loop.** bRT (red) superposed with the group II intron RT GsI-IIC (blue) in ribbon representation. View is to highlight the α-loop of GsI-IIC. Subdomains of the RTs are indicated.
